# GalaxyVS: Exploring 100-Billion Compounds in Seconds

**DOI:** 10.64898/2026.05.29.728912

**Authors:** Xin Hong, Peishun Li, Wenyu Zhu, Chengkun Wu, Hao Guo, Haichuan Tan, Qi Wu, Kelin Wu, Liangsheng Chen, Yinjun Jia, Bowen Gao, Xiaodong Jian, Zhiquan Lai, Youyou Lu, Xiangfei Meng, Yanyan Lan

## Abstract

We present GalaxyVS, a hardware-software co-designed virtual screening framework built to explore the 100-billion commercially accessible chemical space in seconds, deployed at the National Supercomputing Center in Tianjin. Built upon the dense vector retrieval paradigm of DrugCLIP, GalaxyVS bypasses the structural dependencies and computational overhead of classical docking to enable rapid screening against experimentally determined as well as geometrically feasible pockets on AlphaFold-predicted structures. To scale this paradigm to the 100-billion level, the system must overcome the significant computational burden of offline representation encoding, critical memory and I/O bottlenecks during online retrieval, and the risks of diversity collapse and precision loss within final screening results. Utilizing the heterogeneous supercomputing infrastructure, GalaxyVS accelerates the offline encoding through deep operator adaptations and resolves online retrieval bottlenecks via disk-native vector indexing coupled with in-memory staging to ensure both broad accessibility and high throughput. Concurrently, a two-stage refinement protocol effectively mitigates diversity collapse and ensures high-fidelity affinity ranking. Consequently, GalaxyVS achieves a daily scoring throughput of 1.5 × 10^16^ target-ligand pairs, representing a six-orders-of-magnitude leap over previous supercomputing records. Driven by this throughput, we screened nearly 100,000 protein structures across six species against the 100-billion compound library in just 16 hours. The resulting comprehensive cross-species interaction landscape, GalaxyDB, will be openly released at https://galaxyvs.drugclip.com.

## 1 Introduction

Virtual screening serves as a critical step in early-stage drug discovery for identifying structurally novel hit molecules from vast chemical libraries against specific protein targets. However, current screening campaigns have yet to fully leverage the vast biological and chemical space now accessible. Specifically, although AlphaFold2 [1] has predicted structures for 158 million proteins, covering 78% of the 200 million entries in UniProt, this structural repository remains largely untapped even for targets with high therapeutic relevance. Simultaneously, the 100-billion scale of commercially accessible chemical libraries is rarely explored, which restricts the discovery of diverse molecular scaffolds and potent hit molecules [2, 3]. These gaps persist primarily because classical methodologies such as structure-based molecular docking [4–8] and physical simulations like Free Energy Perturbation [9] face critical constraints. First, reliance on high-resolution experimental structures limits the search space to approximately 70,000 proteins archived in the Protein Data Bank, representing a mere 0.035% of known targets. Second, high computational demands restrict scalability, as individual screening campaigns are currently capped at 5.5 billion compounds [10]. Modern drug discovery therefore faces a pressing need for a new screening paradigm capable of simultaneously embracing newly accessible protein targets and the expanding scale of available chemical compounds.

In recent years, deep learning approaches have adopted dense vector retrieval to address these computational and structural constraints, as demonstrated by frameworks like DrugCLIP [11, 12]. By reformulating virtual screening as a maximum inner product search in a shared latent space, this retrieval paradigm improves computational efficiency over traditional physical simulations. Furthermore, pre-training on large-scale synthetic data enables these models to learn robust representations that reduce the reliance on high-fidelity experimental conformations. This structural adaptability supports the direct utilization of AlphaFold2-predicted structures in large-scale screening. Wet-lab experiments have validated this capability by identifying active molecules for TRIP12, a target lacking experimentally resolved structures, using only its predicted pocket. Leveraging these advantages in processing speed and structural adaptability, recent efforts have screened the human proteome against a 500-million compound library and released the resulting dataset to support community research.

The 500-million scale still leaves a significant gap compared to the 100-billion commercially accessible chemical space. However, transitioning to 100-billion-scale exploration introduces three challenges. At the hardware level, generating high-dimensional dense embeddings for 100 billion molecules requires computational and storage capacities that exceed standard computing infrastructures, producing nearly a petabyte of data. At the software level, conventional vector retrieval engines such as FAISS [13] require loading the entire index structure into memory. Even with advanced vector quantization, the memory footprint for a 100-billion vector index imposes extreme resource demands, making traditional in-memory search architectures prohibitively inefficient at such massive scales. At the algorithmic level, maintaining screening fidelity at the 100-billion scale requires addressing the concurrent needs for chemical diversity and ranking precision. Specifically, an unconstrained search across vast libraries often yields results dominated by redundant analogs, while the rapid retrieval paradigm lacks the fine-grained physical modeling essential for accurate affinity prediction.

To bridge these gaps, we present GalaxyVS, a hardware-software co-designed virtual screening framework deployed at the National Supercomputing Center in Tianjin, whose heterogeneous infrastructure directly meets the computational and storage demands of 100-billion-scale processing. By integrating PipeANN [14] for disk-native vector indexing, GalaxyVS enables ultra-large-scale screening on standard nodes with minimal memory overhead. This retrieval engine is further augmented by a distributed partitioning and preloading strategy that asynchronously stages data shards into node-level memory, resolving shared storage I/O bottlenecks under large-scale parallel access. To guarantee screening quality, we introduce a two-stage refinement protocol. First, a diversity control mechanism caps candidate extraction per structural cluster, broadening scaffold coverage across the final hit set. Second, an affinity-based re-ranking stage utilizing AlphaRank [15] explicitly models fine-grained 3D atomic interactions, ensuring high-precision lead prioritization.

Through rigorous in silico evaluations, GalaxyVS demonstrates highly scalable performance. By transitioning binding affinity estimation to a dense retrieval paradigm, the framework exhibits a distinctive sub-linear scaling behavior. The system achieves a daily capacity of 1.5 × 10^16^ target-ligand scorings, effectively shifting the primary computational bottleneck of ultra-large-scale virtual screening (Figure 1). Supported by the disk-native architecture and the two-stage refinement protocol, GalaxyVS successfully mitigates diversity collapse and yields thermodynamically stable candidates with improved binding potential. Leveraging this validated foundation, we deployed GalaxyVS to conduct an extensive virtual screening campaign against the structural proteomes of multiple representative species. In just 16 hours, the system screened nearly 100,000 protein structures against the 100-billion compound library, evaluating over 4 million binding pockets and executing 4.0 × 10^17^ pocket-ligand scorings in total. The resulting cross-species protein-ligand interaction landscape, GalaxyDB, is openly released to support data-driven drug discovery.

**Figure 1.**
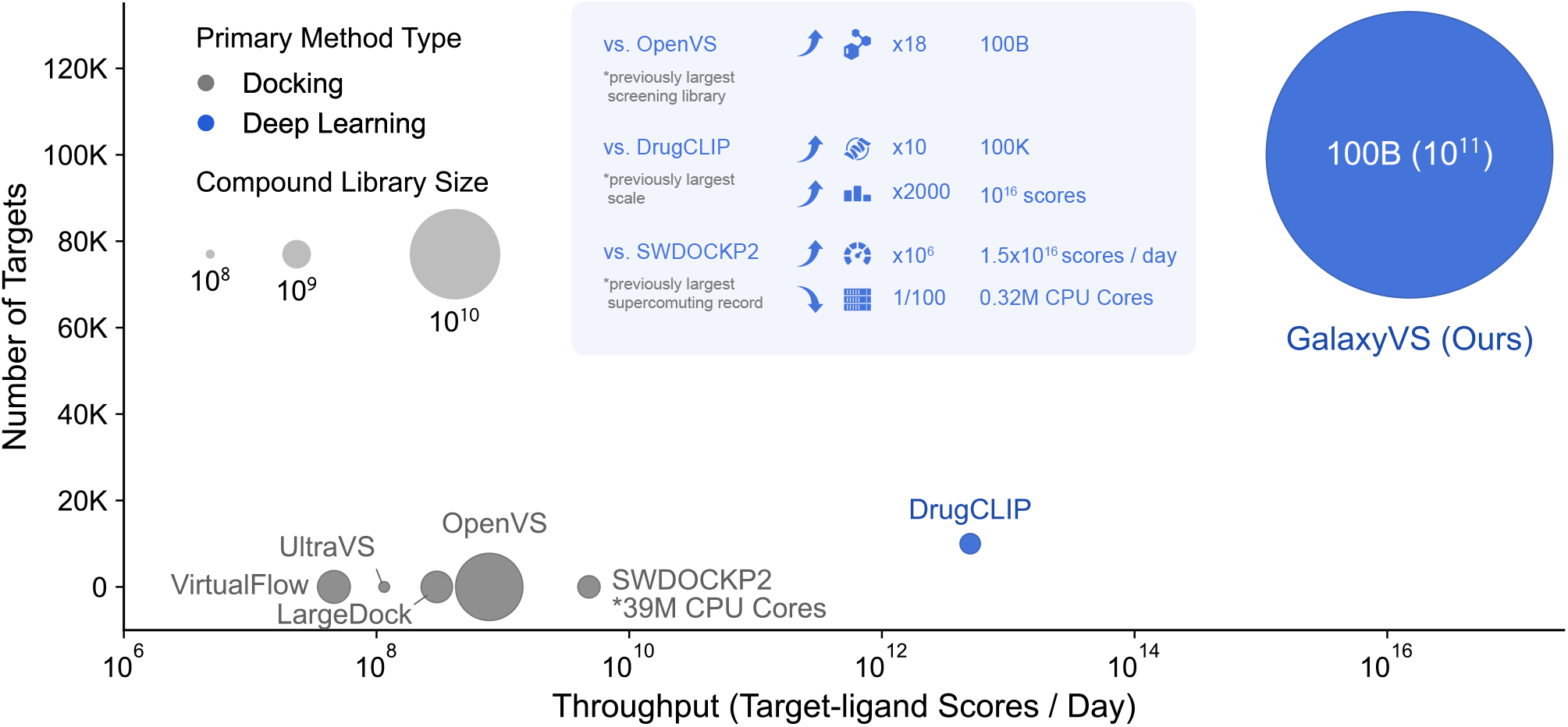
Comparison of GalaxyVS against existing large-scale virtual screening benchmarks. GalaxyVS pushes the frontiers of virtual screening across three critical dimensions. **Library Size**. The platform supports a 100-billion compound library, which represents an 18-fold increase over the previously largest campaign conducted by OpenVS. **Screening Scale**. GalaxyVS systematically explores over 100,000 protein pockets, which provides a 10-fold increase in target coverage and results in 2000 times higher total scorings compared to the previous state-of-the-art results from DrugCLIP. **Computational Efficiency**. The system achieves a daily throughput of 1.5 × 10^16^ scores, which constitutes a million-fold speedup over previous supercomputing records set by SWDOCKP2 while utilizing only 0.32 million CPU cores. These breakthroughs enable the first systematic screening of millions of targets against the 100B-scale chemical space.

## 2 Method

### 2.1 Overview of the GalaxyVS Framework

The emergence of DrugCLIP [11, 12], a multimodal contrastive learning framework that reformulates traditional structure-based virtual screening into a dense vector retrieval task, has provided a feasible roadmap for the rapid screening of ultra-large libraries. By its inherent design, this retrieval-based paradigm decouples the extensive offline library preparation from the online screening process. However, realizing this decoupled vision at a 100-billion scale is not a mere expansion in volume. It confronts substantial barriers in both computational scalability and quality assurance. To address these challenges, we propose GalaxyVS, an ultra-large-scale virtual screening framework illustrated in Figure 2. Rather than a simple integration of existing tools, GalaxyVS is structurally organized to empower this large-scale paradigm from the underlying hardware architecture up to the final affinity-based candidate refinement. The framework systematically executes this pipeline through three interconnected technical pillars.

**Figure 2.**
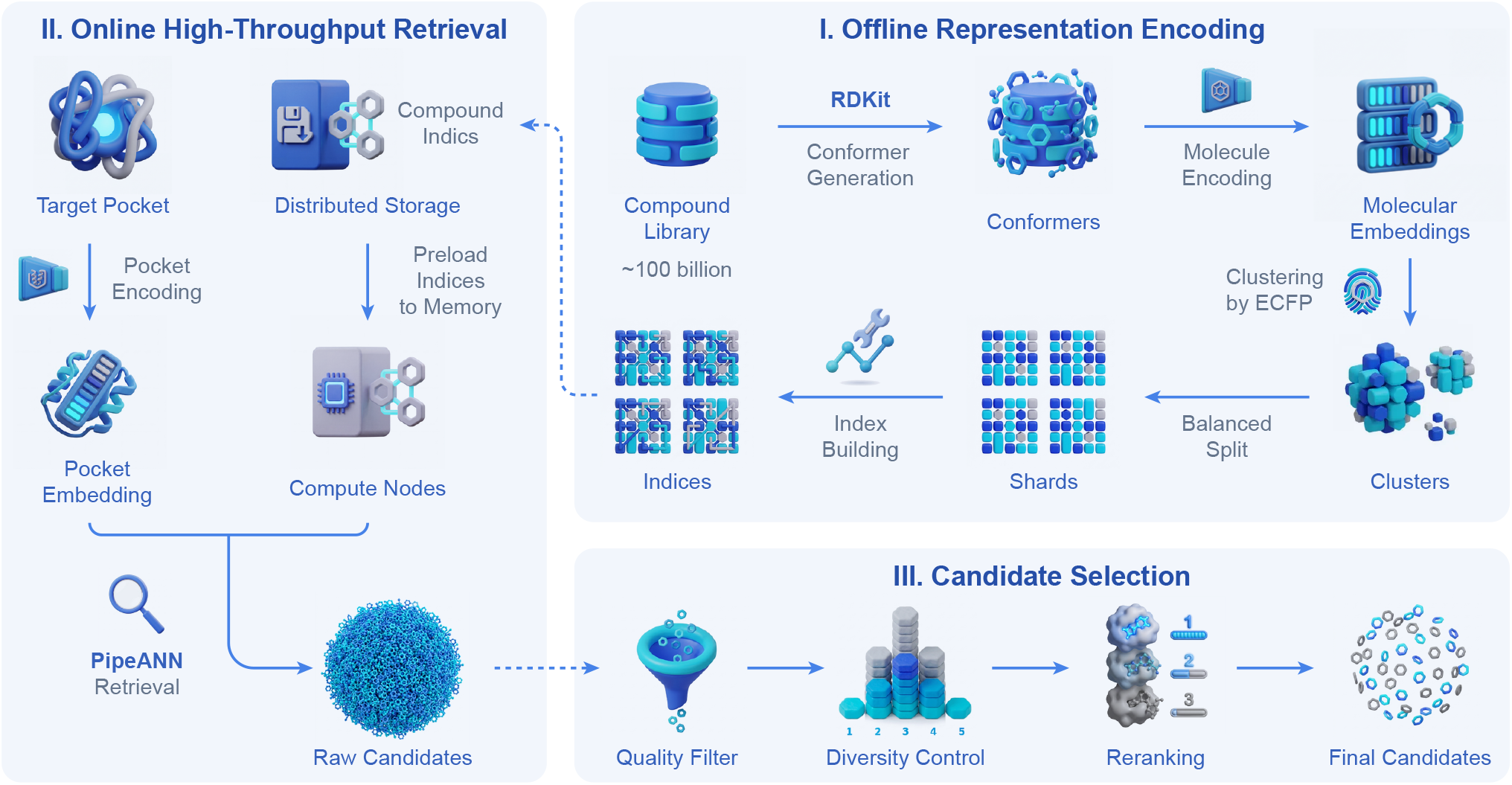
The end-to-end hardware-software co-designed virtual screening pipeline of GalaxyVS. The framework systematically executes ultra-large-scale screening through three interconnected modules. **I. Offline Representation Encoding**. A 100-billion compound library is processed through 3D conformer generation and DrugCLIP encoding to build a static embedding library. To ensure scalability, the library is partitioned into structurally balanced shards based on ECFP-based clustering. **II. Online High-Throughput Retrieval**. Target pockets are encoded into dense embeddings and queried against the distributed indices. This module utilizes PipeANN for disk-native vector search and implements in-memory staging to overcome storage I/O bottlenecks across the supercomputing nodes. **III. Candidate Selection**. The retrieved raw candidates undergo a two-stage refinement protocol. A structural diversity control mechanism first prevents the saturation of dominant scaffolds, followed by an affinity-based re-ranking stage utilizing AlphaRank to explicitly model fine-grained 3D atomic interactions for high-fidelity selection.

**Figure 3.**
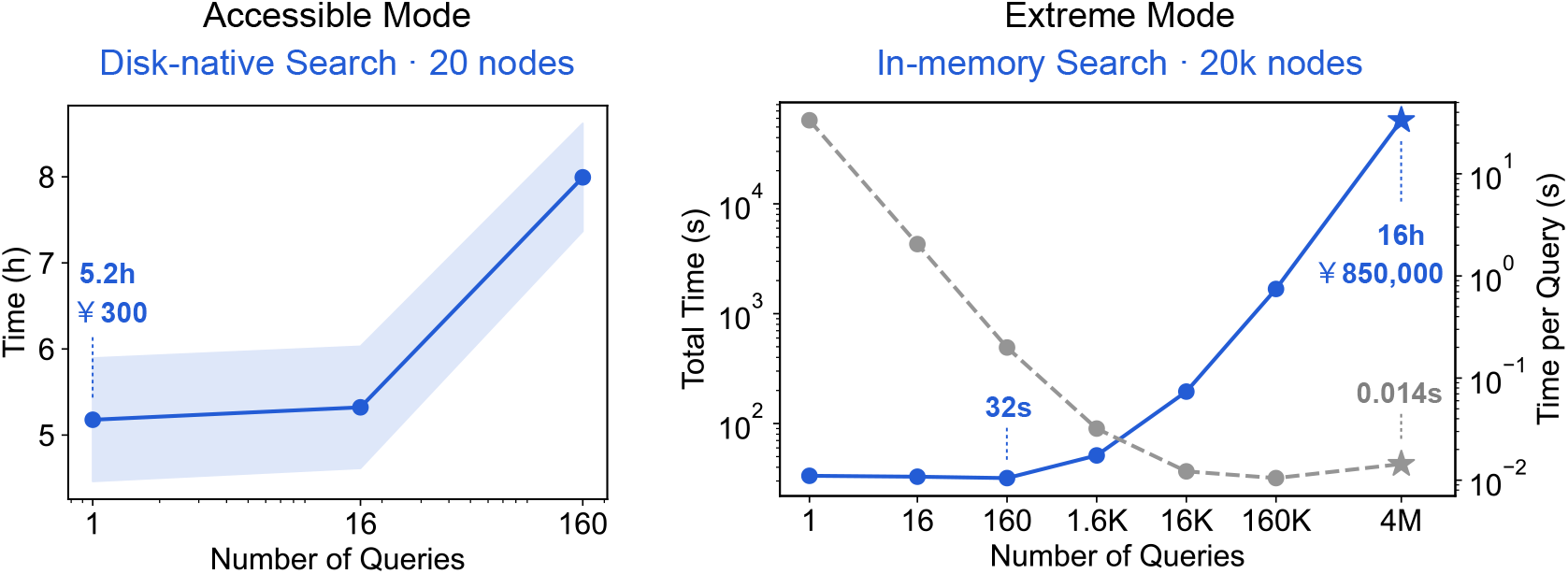
Throughput and cost-efficiency scaling of GalaxyVS across dual operational modes. **Left:** The Accessible Mode (On-disk Search), designed for routine laboratory usage on 20 standard nodes, accomplishes single-target screening against the 100-billion library within 5.2 hours at a highly economical cost (approx. ¥300), enabling practical and budget-friendly ultra-large-scale exploration. **Right:** The Extreme Mode (In-memory Search), deployed across 20,071 computing nodes, demonstrates highly optimized sub-linear scaling, capable of executing millions of concurrent queries (e.g., pan-proteome screening) in just 16 hours.

First, establishing the foundational offline database for 100 billion molecules necessitates the generation of 3D conformations and DrugCLIP-based dense embeddings. This one-time workload demands intensive molecular mechanics optimizations and transformer-based inference for every single molecule, a computational requirement that exceeds the capacity of conventional computing clusters. To address this computational demand, we develop a resilient hybrid task scheduling system and implement deep operator adaptation tailored for heterogeneous architectures. As depicted in the initial phase of the framework, this systematic optimization ensures robust representation encoding across the supercomputing platform to build the static embedding library (Section 2.2).

Second, during the online screening phase, navigating the pre-computed dense embeddings introduces significant storage and bandwidth challenges. To ensure broad accessibility, we integrated PipeANN as a disk-native retrieval engine, which enables ultra-large-scale screening directly from on-disk indices without the need for full memory residency. To support massive parallel discovery, we coupled this engine with a distributed partitioning and preloading strategy to asynchronously preload data shards into node-level memory caches. This architecture effectively overcomes shared storage I/O bottlenecks and achieves extreme throughput for high-concurrency screening tasks (Section 2.3).

Finally, retrieving candidates from a 100-billion-scale chemical space online introduces a diversity collapse dilemma. Because the offline library is vastly expanded, selecting a practical number of top-ranked candidates equates to sampling a minute fraction of the entire pool. Within this narrow top tier, the retrieved molecules tend to be structurally redundant analogs tightly clustered in the semantic space. Furthermore, while DrugCLIP’s decoupled dual-encoder architecture accelerates virtual screening by efficiently enriching active molecules, its design purposely trades fine-grained physical modeling for unprecedented speed. To guarantee the highest quality of the final selection, we resolve these challenges through a unified two-stage candidate refinement protocol. We first introduce a structural diversity control mechanism to enforce chemical novelty across the pool, followed by an affinity-based re-ranking stage utilizing AlphaRank [15] to seamlessly complement the retrieval’s architectural trade-off. Ultimately, this comprehensive refinement protocol ensures that the rapid vector retrieval yields a candidate pool that is both chemically distinct and enriched for compounds with high predicted activity. (Section 2.4).

### 2.2 Large-Scale Representation Encoding on Heterogeneous Architectures

The theoretical foundation of our virtual screening pipeline relies on DrugCLIP [11, 12], a multimodal contrastive learning framework that fundamentally breaks the traditional trade-off between screening throughput and predictive accuracy. Unlike conventional structure-based docking that requires computationally expensive pose sampling and scoring for every protein-ligand pair, DrugCLIP employs two distinct Uni-Mol-based transformer encoders. Through contrastive learning, it implicitly captures complex binding patterns and aligns the spatial features of both modalities into a shared representation space. This paradigm shift reformulates the geometry-dependent affinity estimation into a highly efficient cosine similarity calculation between dense vectors. Crucially, this dual-encoder architecture fully decouples the ligand encoding process from the target protein, providing the structural prerequisite to pre-compute and store representations for the entire chemical library.

Leveraging this decoupled paradigm, we construct the massive offline database for 100 billion molecules. The data processing pipeline begins by converting SMILES strings into 3D molecular conformations using RDKit [16]. These generated geometries are then systematically fed into the DrugCLIP molecule encoder to extract static representation vectors.

Executing this inference pipeline at a 100-billion scale demands immense computational throughput and robust system-level orchestration. To ensure optimal scalability and accommodate the model’s operational requirements, we adapted the encoding workflow to the heterogeneous computing architecture of the Tianhe supercomputer. Utilizing the YH-Torch intelligent computing framework, we performed deep operator-level adaptations to optimize the memory hierarchy and minimize data transfer overheads on the AI accelerators. Concurrently, to manage the massive task concurrency, we deployed a resilient scheduling system equipped with automated fault-tolerance mechanisms. This hardware-software co-optimization guarantees the stable progression of the encoding workflow. Finally, to enhance the robustness of the generated representations against conformational variances, the embedding for each molecule is aggregated from an ensemble of six independently trained DrugCLIP models, yielding a reliable static library ready for downstream retrieval.

### 2.3 Efficient High-Throughput Vector Retrieval

Navigating a 100-billion-scale embedding library introduces substantial storage and bandwidth challenges. While conventional in-memory vector search frameworks such as FAISS provide highly efficient retrieval, maintaining full index residency in memory at this scale requires large aggregate RAM capacity, which amounts to hundreds of terabytes, across the computing cluster. To improve accessibility under resource-constrained settings, we integrated PipeANN as a disk-native retrieval engine that supports graph search directly over on-disk indices without requiring complete memory residency. This capability provides the technical foundation for the *Accessible Mode* of GalaxyVS, enabling ultra-large-scale screening on a limited number of standard compute nodes with modest memory requirements.

To further support massive parallel tasks such as pan-proteome screening, we extended this architecture into a high-concurrency *Extreme Mode*. While the disk-native approach ensures accessibility, direct expansion of node counts can lead to severe I/O congestion on shared file systems. Inspired by the high performance of in-memory search, we implemented a partitioning and preloading strategy. The 100-billion library is divided into structurally balanced shards that are asynchronously preloaded from the shared storage into the local memory of compute nodes. This strategy effectively decouples search execution from shared disk I/O bottlenecks. Once loaded, the system executes rapid parallel retrieval directly from the in-memory data, enabling 100-billion-scale screening for batches of pockets in seconds.

### 2.4 Candidate Selection via Structural Diversity Control and Affinity Re-ranking

Following the initial vector retrieval, the raw cosine similarity scores across multiple target pocket conformations are first calibrated using an adjusted robust Z-score normalization. This calibration is a fundamental step adopted from the DrugCLIP [11, 12] paradigm to ensure score comparability. While this high-throughput retrieval efficiently narrows down the 100-billion library, it introduces two critical challenges at this scale. First, a naive score-based truncation of the top candidates inevitably leads to diversity collapse, where the resulting pool is overwhelmingly saturated with minor structural variations of a few dominant scaffolds. Second, the decoupled dual-encoder architecture of DrugCLIP prioritizes retrieval speed and broad enrichment over fine-grained physical interaction modeling. To ensure the chemical novelty and biochemical relevance of the final selection, we implement a comprehensive candidate refinement protocol.

To overcome diversity collapse, we designed a structural diversity control mechanism built upon the structurally-aware partitioning introduced in Section 2.3. Prior to index construction, the entire molecular library is clustered into approximately 10,000 structural families using K-Means clustering based on ECFP4 fingerprints. Because this clustering inherently yields highly unbalanced cluster sizes, we uniformly subdivide them to resolve the imbalance and reduce the computational overhead per retrieval task, ultimately forming 100,000 structurally cohesive data shards of roughly equal size. During the candidate selection phase, we introduce a dynamic diversity factor to regulate the retrieval distribution. By strictly capping the maximum number of high-scoring molecules contributed by any single cluster, this factor forces the final selection to be sourced from a much broader range of clusters. This strategy successfully elevates the overall chemical diversity while preserving the highest possible screening quality.

As a critical refinement to further elevate the virtual screening performance, the structurally diverse candidate set is subsequently processed through an affinity-based re-ranking stage. While DrugCLIP provides highly effective and rapid initial enrichment, its architecture purposely trades fine-grained interaction modeling for unprecedented speed. To seamlessly complement this trade-off, we employ AlphaRank [15], a sophisticated interaction model featuring a protein-ligand co-folding architecture. Unlike the coarse-grained semantic embeddings used in the retrieval phase, AlphaRank explicitly models the 3D atomic-level interactions and spatial conformations between the target pocket and the ligand. This final step re-evaluates the candidates based on high-fidelity predicted interaction strengths, successfully refining mathematically enriched nearest neighbors into biochemically rigorous drug candidates.

## 3 Results

### 3.1 GalaxyVS Enables 100-billion-scale Virtual Screening in Seconds

The dual-mode architecture of GalaxyVS provides both routine accessibility and extreme throughput for ultra-large-scale virtual screening. For regular laboratory usage, GalaxyVS provides an *Accessible Mode* that leverages the disk-native retrieval capability of PipeANN. This mode enables a single-target screen against the 100-billion compound library using only 20 compute nodes in approximately 5.2 hours. With an estimated cost of 300 CNY per target, this approach ensures that ultra-large-scale exploration is economically viable for individual research projects without requiring specialized high-performance hardware.

While the disk-native approach ensures accessibility, the discovery of cross-species proteome-wide interactions demands a much higher level of computational intensity. To overcome the I/O bottlenecks inherent in shared storage, we implemented a distributed partitioning and preloading architecture for the *Extreme Mode*. By pre-caching index shards into node-level memory caches, the framework effectively transitions disk-bound retrieval into high-speed local searches. Under this configuration, GalaxyVS completes a single screening run for a batch of 160 binding pockets against the 100-billion library in just 32 seconds. When GalaxyVS operates at full power to conduct exhaustive screening campaigns, it achieves a record-breaking daily throughput of 1.5 × 10^16^ scorings. This performance represents a million-fold increase over previous supercomputing records for docking-based virtual screening.

The breakthrough in throughput is enabled by a paradigm shift that moves the primary computational burden to an offline data preparation phase. Resource-intensive procedures such as 3D conformer generation and molecular embedding are treated as one-time sunk costs. Constructing the 100-billion-scale library required 1,000 CPU nodes for 12 days and 5,000 heterogeneous accelerator nodes for 18 days. GalaxyVS bypasses these overheads during online retrieval by converting binding predictions into near-instantaneous vector operations, achieving minimal runtime latency through a high-concurrency memory-preloading architecture. Because high-concurrency vector retrieval remains highly efficient even as batch sizes grow, the effective per-pocket latency approaches approximately 0.014 seconds when scaling to millions of concurrent queries. This sub-linear scaling behavior effectively shifts the primary computational bottleneck of drug discovery from years of simulation to seconds of retrieval.

### 3.2 GalaxyVS Delivers Diverse and Novel Hits

We quantified the structural diversity of GalaxyVS hits across 102 DUD-E targets and compared them against a representative in-stock screening library aggregated from ChemDiv, ChemBridge, Enamine, and Life Chemicals. After apply physicochemical and structural filters, the baseline library contained 2.94M compounds. For each target, the top 30,000 compounds (∼ 1%) ranked by score were retained from the baseline library. For the ultra-large library, the top 0.01% retrieved compounds were subjected to the same filtering procedure, followed by diversity-controlled selection to preserve 30,000 compounds spanning approximately 8,000 library clusters.

To systematically characterize chemical diversity, we performed strict leader-clustering on each candidate pool and quantified the numbers of unique scaffolds and molecular fragments. As illustrated in Figure 4, GalaxyVS consistently recovers a broader distribution of structurally distinct clusters, while yielding substantially higher numbers of unique scaffolds and fragments compared with the conventional million-scale library. These results suggest that GalaxyVS effectively alleviates the structural redundancy commonly observed in conventional virtual screening pipelines.

**Figure 4.**
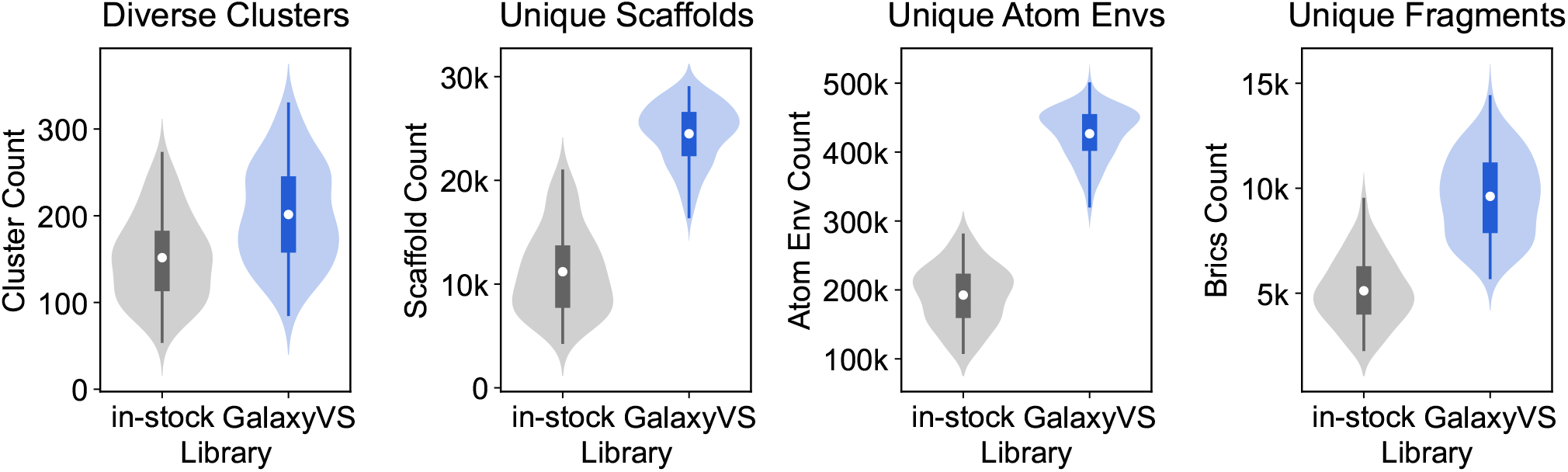
Pairwise diversity comparison of top-ranked 30,000 compounds across 102 DUD-E targets. GalaxyVS hits retrieved from the ultra-large library yields more highly diverse structural clusters, as well as a higher number of unique scaffolds and fragments, compared with a representative million-level screening library.

Beyond structural diversity, we further investigated the translational potential of the retrieved hits by evaluating their chemical novelty within the clustered candidate pools. As illustrated in Figure 5, the candidate sets derived from the in-stock library and GalaxyVS exhibit different novelty profiles across 42 targets with novelty challenges in either screening setting. The circular bar charts categorize chemical novelty into four levels, reflecting the degree of overlap with known chemical space and potential intellectual property constraints.

**Figure 5.**
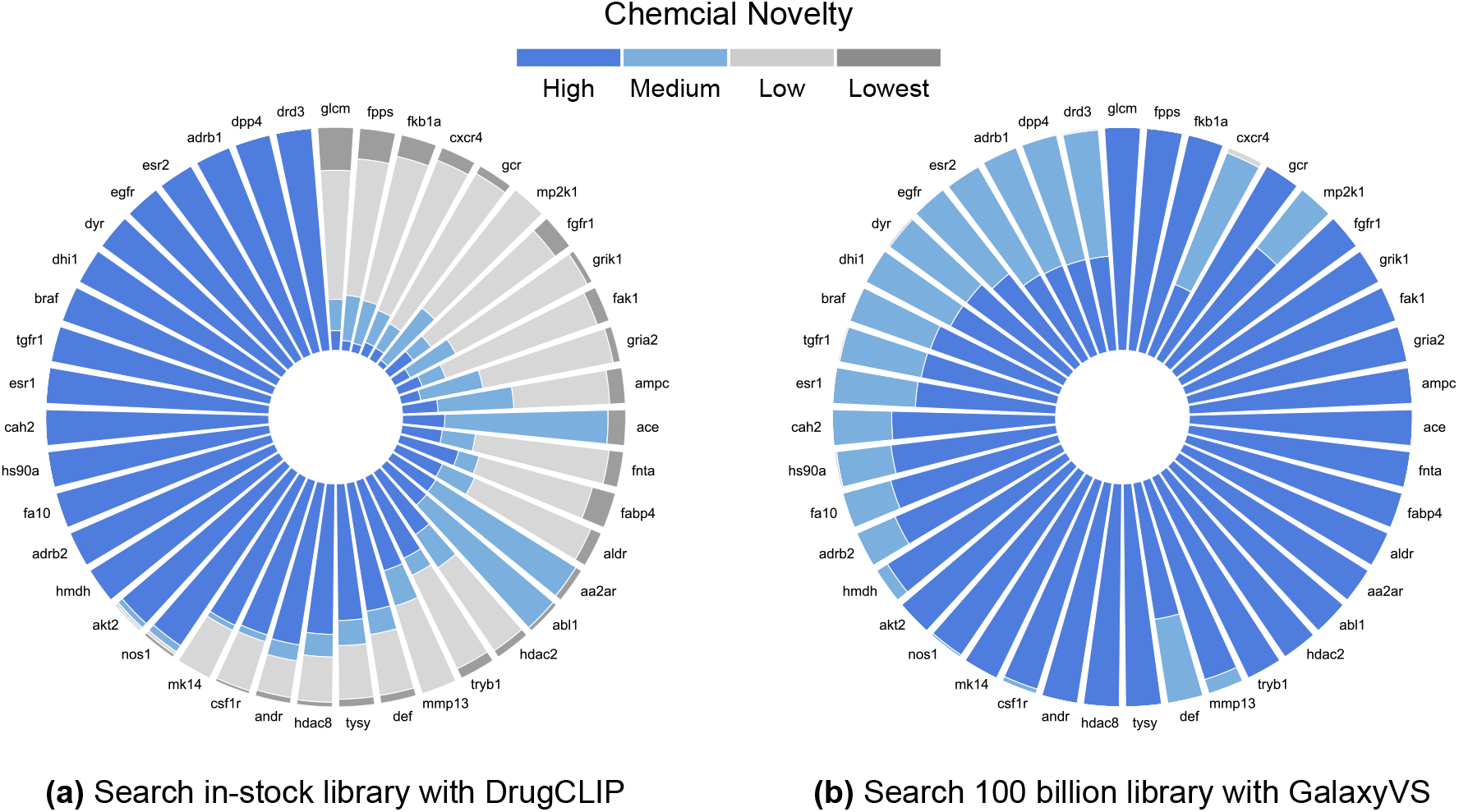
Chemical novelty profiles of clustered hit pools across 42 targets exhibiting novelty challenges in either the in-stock or ultra-large screening libraries. Colors blue, light blue, light gray, and gray represent decreasing levels of chemical novelty, ranging from high to the lowest. Based on PubChem search, these four categories correspond to (i) highly novel compounds (unrecorded with *<* 5 analogs), (ii) medium novelty structures (either recorded with *<* 5 analogs or unrecorded with ≥ 5 analogs), (iii) low novelty clusters (recorded with ≥ 5 analogs), and (iv) preexisting patent-covered compounds, respectively. Here, analogs are defined by PubChem fingerprints with the default similarity threshold at 0.9.

The comparison reveals that conventional virtual screening against the in-stock library produces a substantially larger proportion of compounds within low and the lowest novel categories across targets, indicating that the enriched hits are concentrated in highly crowded and potentially patent-constrained chemical regions. In contrast, GalaxyVS demonstrates a significantly improved novelty profile with access to substantially less explored chemical space. Even for the most challenging targets, the retrieved compounds are predominantly distributed within the medium-novelty regime, corresponding to previously unreported compounds that retain only partial substructure similarity to known molecules. This observation suggests that GalaxyVS can effectively move beyond heavily saturated chemical series while preserving favorable retrieval quality.

In summary, these results demonstrate that GalaxyVS not only expands structural diversity, but also substantially improves access to novel and less crowded chemical space. Such broad and chemically distinct coverage facilitates both the discovery of novel hit scaffolds and the identification of multiple optimization chemical series for downstream research and development.

### 3.3 GalaxyVS Enriches High-affinity Predicted Binders

While structural diversity and chemical novelty broaden the exploring space, the value of the compounds primarily depend on their binding affinity to targets of interest. At a 0.01% retrieval depth over the 100-billion compound library, GalaxyVS preserves DrugCLIP’s enrichment performance while exposing a substantially larger pool of potential active compounds in the top-ranked set. Provided that the overall positive rate of the library does not decrease by orders of magnitude, this expansion in search space is expected to yield candidates with improved binding affinity at comparable selection sizes. To evaluate this hypothesis, Boltz-2 [17] was applied as an affinity predictor to assess the binding quality of diverse candidates retrieved for each target. For both the ultra-large library and the in-stock screening library, we selected up to 60 representative compounds per target using cluster leaders to control for chemical redundancy and approximate a realistic experimental budget.

The Boltz-2 score distribution for the 100B library is clearly shifted toward stronger predicted affinities relative to the million-level baseline (Figure 6a), and this shift is consistent across most evaluated targets (Figure 6c). Aggregated by protein class (Figure 6b), GalaxyVS achieves an overall win rate of 80.4% across the 102 diverse targets. The trend is consistent across the better-represented classes, with win rates above 70% for enzymes (kinases 76.9%, *n*=26, proteases 93.3%, *n*=15 and others 83.3%, *n*=36) and nuclear receptors (72.7%, *n*=11); the smaller subsets (ion channels, P450, miscellaneous; *n* ≤ 5) follow the same direction but are too sparse to support per-class conclusions. The one apparent deviation is GPCRs (40.0%, *n*=5); although this estimate is also limited by sample size, the result is consistent with the high conformational flexibility of these receptors, where static scoring functions struggle in distinguishing true binders from background library noise.

**Figure 6.**
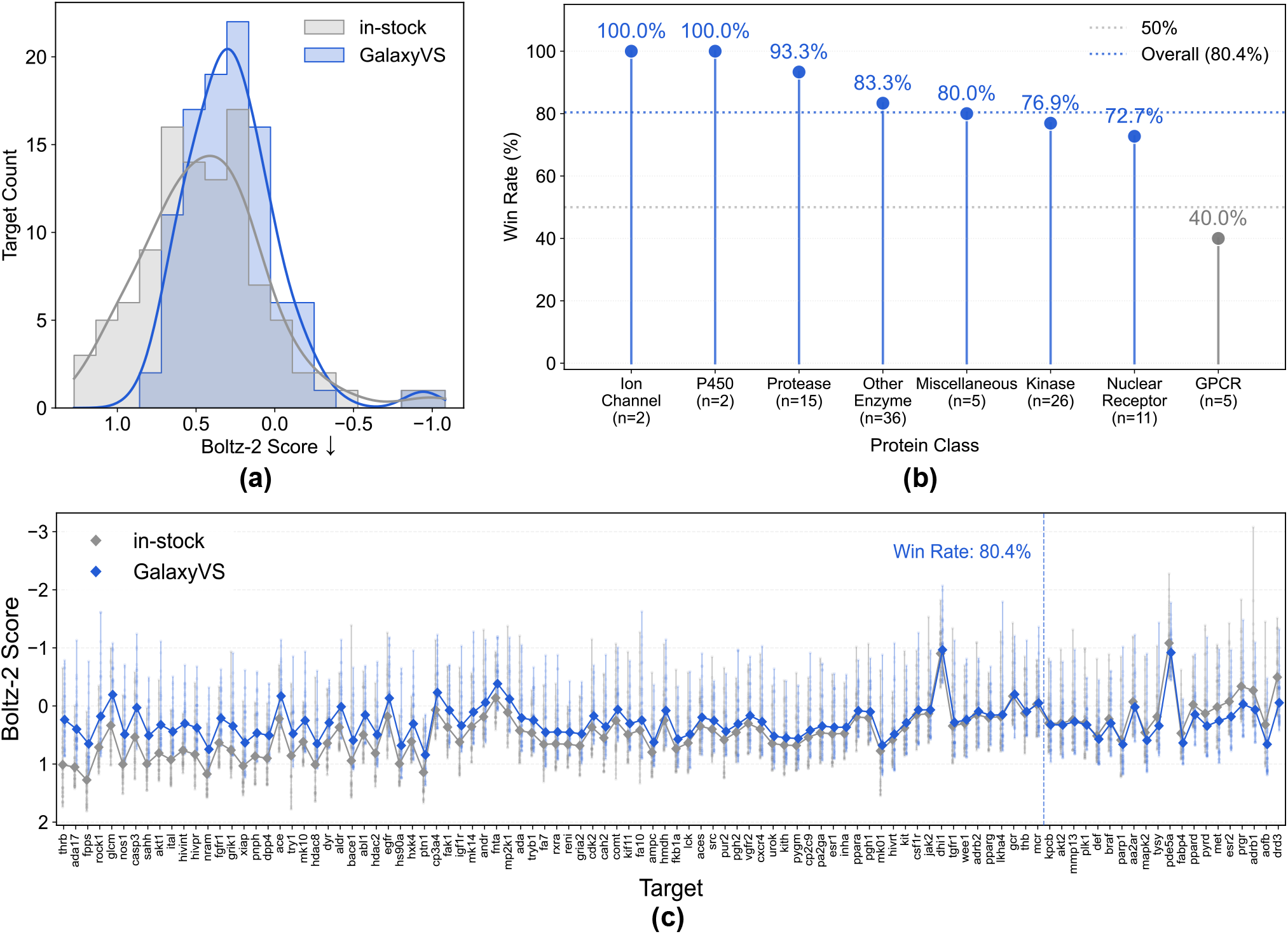
Performance of GalaxyVS across various protein classes (102 DUD-E targets) in term of predicted affinity. **(a)** Distribution of per-target Boltz-2 scores for compounds from the in-stock library (gray) versus the 100B ultra-large library (blue); more negative scores indicate stronger predicted affinities. **(b)** Win-rate by protein class, defined as the fraction of targets for which GalaxyVS achieve a better mean Boltz-2 score than the in-stock baseline; the dotted line marks the overall win-rate (80.4%). **(c)** Per-target Boltz-2 score distributions of the top 60 retrieved compounds in each library.

**Figure 7.**
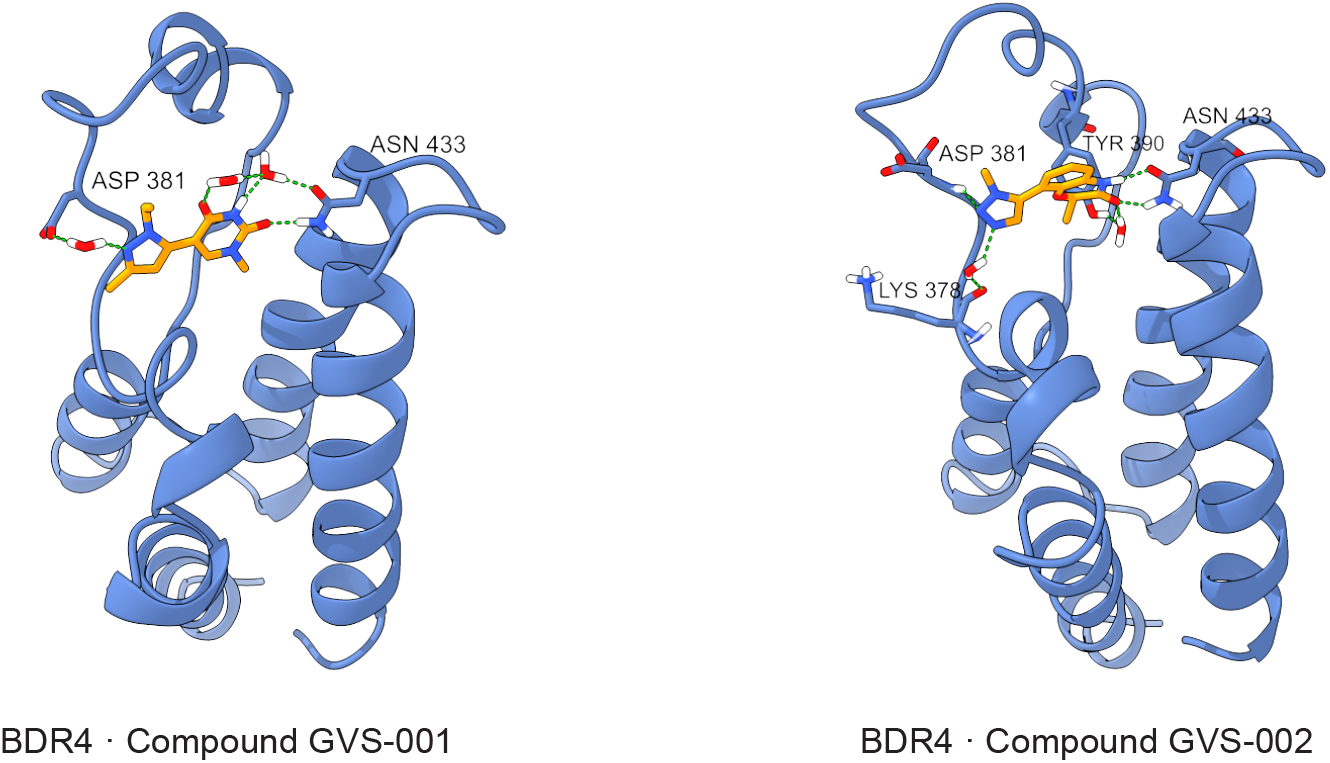
Equilibrated binding modes of the two top-scored compounds in the ABFEP simulations. Among the ten candidates screened against BRD4 (PDB ID: 5UF0), compounds GVS-001 and GVS-002 exhibit Δ*G* values of −8.97 *±* 0.89 kcal/mol and −8.03 *±* 0.85 kcal/mol, respectively.

To further validate the enrichment capability of GalaxyVS from a physics-based perspective, we performed Absolute Binding Free Energy Perturbation (ABFEP) [18] calculations using BAT.py [19]. From a filtered and clustered pool, the top 10 molecules were subjected to molecular dynamics and thermodynamic evaluation against BRD4. The selected candidates exhibit consistent binding affinities, achieving an average Δ*G* of -6.25 kcal/mol and a standard deviation of 1.57 kcal/mol. The calculations yielded reliable results with an average statistical error of 1.12 kcal/mol, highlighted by favorable candidates such as Compound GVS-001 (Δ*G* = -8.97 ± 0.89 kcal/mol) and Compound GVS-002 (Δ*G* = -8.03 ± 0.85 kcal/mol), indicating robust and well-converged thermodynamic behavior.

To understand the structural basis of this consistency, we analyzed representative binding poses from the ABFEP simulations. Compounds GVS-001 and GVS-002 adopt canonical binding modes featuring polar interactions with ASP381 and the key anchoring residue ASN433, while forming stable interaction networks with conserved water molecules and the protein backbone. This combination of thermodynamic consistency and well-defined binding modes suggests that GalaxyVS effectively prioritizes ligands that establish persistent and physically meaningful interactions, providing a reliable basis for subsequent experimental validation.

Taken together, these results indicate that GalaxyVS consistently prioritizes compounds with higher predicted and physics-resolved binding affinity across diverse targets, supported by both scoring-based evaluation and free-energy-based validation.

### 3.4 GalaxyVS Explores Proteome-wide Interactions across Multiple Species

The combination of high computational throughput and structural adaptability enables virtual screening to expand from individual targets to comprehensive proteome-level exploration. Leveraging these technical foundations, we deployed GalaxyVS to perform extensive screening against the structural proteomes of multiple representative species using the 100-billion-molecule library. This systematic mapping across biological domains allows for the large-scale identification of protein-ligand interactions, providing a data-driven resource to support diverse research directions such as human disease treatment, antimicrobial discovery, and the development of pesticides or herbicides.

We evaluated the system by processing six representative species that span distinct evolutionary branches. These organisms include *Escherichia coli, Saccharomyces cerevisiae, Arabidopsis thaliana, Drosophila melanogaster, Mus musculus*, and *Homo sapiens*. Structural data were sourced from the AlphaFold Protein Structure Database to extract complete structural proteomes. Following the identification of valid binding sites, the screening space comprised 4 million binding conformers derived from approximately 100,000 protein targets as detailed in Appendix Table 7. This selection covers biological diversity across microbial pathogens and human therapeutic targets.

The cross-species campaign was executed across 20,071 compute nodes within 16 hours. We utilized the hierarchical distinction between target-ligand scorings and pocket-ligand scorings to characterize the computational workload. While target-level evaluation involves the aggregation of results from multiple binding sites for each target, pocket-ligand scoring represents the fundamental retrieval operation between a specific pocket embedding and a molecule. The exhaustive screening of 4 million pockets against the 100-billion-compound library resulted in a total of 4.0 × 10^17^ pocket-ligand scorings. The system maintained an effective search latency of approximately 0.01 seconds per pocket. These results provide a foundation for data-driven discovery. The resulting interaction landscape is released as GalaxyDB to support research in drug discovery.

## 4 Conclusion

Empirical evaluations demonstrate that GalaxyVS delivers performance across two complementary operational modes. The Accessible Mode enables single-target screening against the 100-billion library within 5.2 hours on 20 standard nodes at an estimated cost of 300 CNY per target, making ultra-large-scale exploration economically viable for routine laboratory use. The Extreme Mode exploits sub-linear scaling behavior to complete a batch of 160 pocket queries in just 32 seconds, sustaining a daily throughput of 1.5 × 10^16^ target-ligand pairs and representing a million-fold increase over previous supercomputing records. Our diversity control mechanism effectively mitigates structural redundancy, yielding a broader distribution of unique molecular scaffolds compared to conventional million-scale libraries. Evaluation using the Boltz-2 scoring function demonstrates an overall win-rate of 80.4% over commercial in-stock baselines across 102 DUD-E targets. Further validation via Absolute Binding Free Energy Perturbation (ABFEP) simulations confirms that the identified hits exhibit robust thermodynamic stability. Leveraging its scalable architecture, GalaxyVS executed 4.0 × 10^17^ pocket-ligand scorings in 16 hours across six representative species. This extensive campaign provides a comprehensive cross-species interaction landscape, GalaxyDB, established as an open resource to accelerate data-driven drug discovery. Future work will focus on wet-lab experimental validations to further explore the therapeutic potential of the identified chemotypes.

## Supporting information

Comparison of Large-Scale Clustering Methods

## Appendix

### A Related Works

#### A.1 Compound Libraries for Virtual Screening

The concept of chemical space encompasses an estimated 10^60^ possible drug like molecules [20], providing the theoretical framework for exploring bioactive compounds. Historically, enumerated libraries, such as the widely used ZINC database [21, 22], explicitly list each molecular structure. The field has since evolved toward combinatorial libraries, which are defined by sets of building blocks and reaction rules. This shift enables a combinatorial explosion, efficiently representing vast, synthetically accessible chemical regions and facilitating ultra large libraries, including ZINC22, Enamine REAL Space, WuXi GalaXi, and expanded public collections—that contain billions to trillions of virtual compounds. Key features of these prominent libraries are summarized in Table. Our work utilizes fully enumerated subsets from Enamine REAL Space and WuXi GalaXi.

#### A.2 Recent Large-Scale Virtual Screening Campaigns

Over the past decade, the rapid expansion of compound libraries and the advent of combinatorial chemistry have driven the evolution of ultra-large-scale virtual screening. This section summarizes representative efforts in the field as outlined in Table 2. Technically, docking-based methods [2, 10, 23, 25, 26] dominate this landscape, primarily accelerated through the application of active learning to prioritize promising subsets and the improvement of parallel computing efficiency via hardware-software co-design. In contrast, freeenergy perturbation approaches are rarely employed for large-scale screening due to their high computational cost, with their urgent application during the COVID-19 pandemic to approved drug libraries serving as a notable exception [24]. Recently, deep learning-based virtual screening has emerged as a transformative alternative. For instance, DrugCLIP [12] achieved a breakthrough in throughput by screening nearly 10,000 human proteins against 500 million compounds in approximately one day using only eight GPUs. Despite these advances, current screening campaigns have not yet exceeded tens of billions of compounds, leaving the 100-billion-scale chemical space accessible in commercial libraries largely unexplored. Consequently, the development and application of virtual screening technology must continue to accelerate to keep pace with the exponential growth of available compound libraries.

**Table 1.** Comparison of Common Compound Libraries. *Note:* Library sizes update dynamically. Data retrieved from BioSolveIT chemical spaces catalogue and individual vendor websites, accessed May 2026.

| Library Name | Provider | Size | Feasibility Rate | Delivery Time |
| --- | --- | --- | --- | --- |
| Enamine REAL Space | Enamine | ~95 Billion | > 80% | 3–4 Weeks |
| Enamine xREAL Space | Enamine | >4.4 Trillion | > 80% | 3–4 Weeks |
| WuXi GalaXi | WuXi AppTec | >25.8 Billion | 60–80% | 4–8 Weeks |
| Freedom Space | Chemspace | 296 Billion | > 80% | 5–6 Weeks |
| eXplore | eMolecules | 5 Trillion | > 85% | 3–4 Weeks |
| Synple Space | Synple Chem | 1 Trillion | > 70% | 3–4 Weeks |
| PharmaBlock Sky Space | PharmaBlock | 58 Billion | > 85% | 4–6 Weeks |
| ZINC22 | Public (UCSF) | >37 Billion | Variable | Variable |
| GDB-17 | Academic | 166 Billion | Theoretical | N/A |
| LifeCheMyriads | Life Chemicals | 50 Billion | High | 4–6 Weeks |
| D2B-SpaceM1 | Molecule.One | 1.5 Billion | > 85% | 2–6 Weeks |
| AMBrosia | Ambinter | 126 Billion | High | 4–6 Weeks |
| VAST™ Space | XtalPi | 4.66 Billion | > 85% | 2–4 Weeks |
| KnowledgeSpace | BioSolveIT | 260 Trillion | N/A | N/A |
| CHEMriya | Otava | 55 Billion | High | 4–6 Weeks |

**Table 2.**
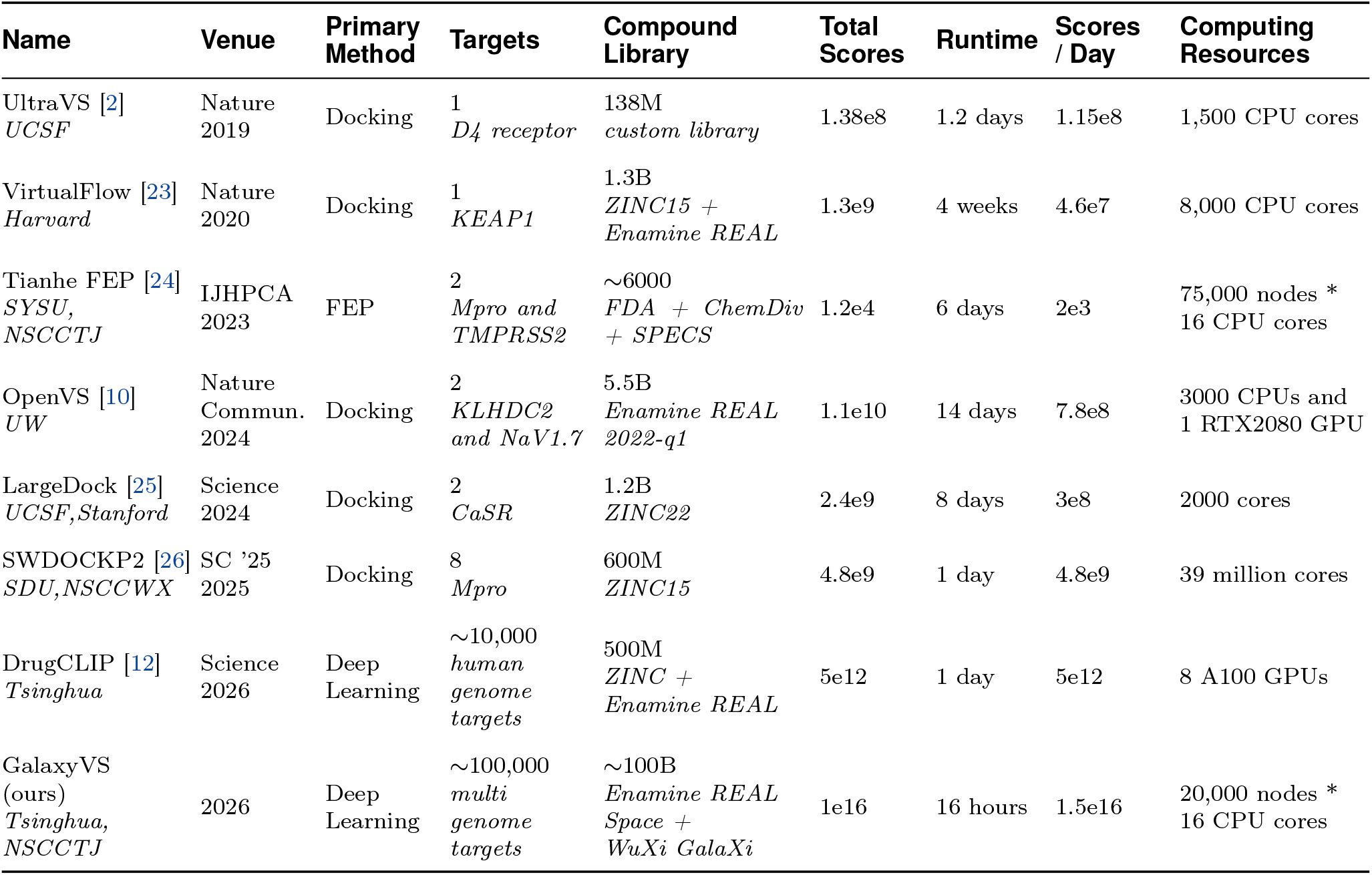
Comparison of Large-scale Virtual Screening Platforms. A single score is defined as one complete evaluation of a protein–ligand pair, including those across multiple pockets and conformations.

| Name | Venue | Primary Method | Targets | Compound Library | Total Scores | Runtime | Scores / Day | Computing Resources |
| --- | --- | --- | --- | --- | --- | --- | --- | --- |
| UltraVS [2]<br>*UCSF* | Nature 2019 | Docking | 1<br>*D4 receptor* | 138M<br>*custom library* | 1.38e8 | 1.2 days | 1.15e8 | 1,500 CPU cores |
| VirtualFlow [23]<br>*Harvard* | Nature 2020 | Docking | 1<br>*KEAP1* | 1.3B<br>*ZINC15 + Enamine REAL* | 1.3e9 | 4 weeks | 4.6e7 | 8,000 CPU cores |
| Tianhe FEP [24]<br>*SYSU, NSCCTJ* | IJHPCA 2023 | FEP | 2<br>*Mpro and TMPRSS2* | ~6000<br>*FDA + ChemDiv + SPECS* | 1.2e4 | 6 days | 2e3 | 75,000 nodes *<br>16 CPU cores |
| OpenVS [10]<br>*UW* | Nature Commun. 2024 | Docking | 2<br>*KLHDC2 and NaV1.7* | 5.5B<br>*Enamine REAL 2022-q1* | 1.1e10 | 14 days | 7.8e8 | 3000 CPUs and<br>1 RTX2080 GPU |
| LargeDock [25]<br>*UCSF, Stanford* | Science 2024 | Docking | 2<br>*CaSR* | 1.2B<br>*ZINC22* | 2.4e9 | 8 days | 3e8 | 2000 cores |
| SWDOCKP2 [26]<br>*SDU, NSCCWX* | SC '25 2025 | Docking | 8<br>*Mpro* | 600M<br>*ZINC15* | 4.8e9 | 1 day | 4.8e9 | 39 million cores |
| DrugCLIP [12]<br>*Tsinghua* | Science 2026 | Deep Learning | ~10,000<br>*human genome targets* | 500M<br>*ZINC + Enamine REAL* | 5e12 | 1 day | 5e12 | 8 A100 GPUs |
| GalaxyVS (ours)<br>*Tsinghua, NSCCTJ* | 2026 | Deep Learning | ~100,000<br>*multi genome targets* | ~100B<br>*Enamine REAL Space + WuXi GalaXi* | 1e16 | 16 hours | 1.5e16 | 20,000 nodes *<br>16 CPU cores |

**Table 3.**
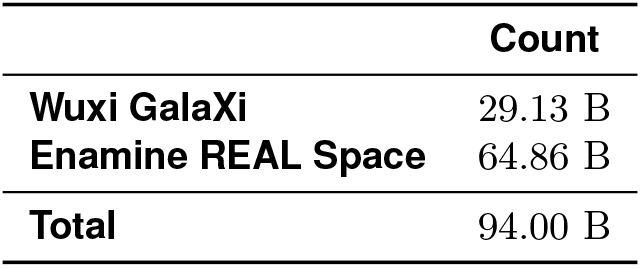
Summary of the compound libraries used in GalaxyVS.

### B Data Curation and Preprocessing

#### B.1 Ultra-Large Compound Libraries

##### Enamine REAL Space

The Enamine REAL Space database represents a carefully curated collection of synthetically accessible small molecules derived from 181,288 qualified building blocks and reagents available from Enamine’s in-stock inventory. These compounds are generated through 172 well-validated synthetic protocols that encompass both standard and advanced one-pot reactions, with variations in synthesis steps, purification methods, and handling requirements to ensure practical feasibility. The library is characterized by its emphasis on synthetic accessibility, with all designed compounds requiring no more than three synthetic steps from available starting materials, resulting in an average synthesis success rate exceeding 80% and a reliable delivery timeframe of 3-4 weeks. This collection demonstrates substantial structural diversity while maintaining strong drug-like properties through careful design parameters. The July 2024 version used in this study contains approximately 64.9 billion compounds, providing extensive coverage of synthetically feasible chemical space for virtual screening applications.

##### Wuxi Galaxi Library

The Wuxi GalaXi virtual compound library, provided by WuXi AppTec through LabNetwork, serves as a fundamental screening resource within the GalaxyScreen platform. This computationally enumerated collection supports early-stage drug discovery by integrating over 30 parallelizable reaction types and leveraging WuXi’s proprietary repository of building blocks, scaffolds, and templates that have been accumulated through more than 20 years of medicinal chemistry expertise. Each compound is optimized for drug-likeness using key physicochemical parameters including molecular weight, LogP, TPSA, and ADMET properties, while being constrained to a maximum of three synthetic steps from in-stock reagents to ensure high synthetic feasibility rates of 60 to 80 percent. The library was strategically selected for GalaxyScreen’s trillion-scale virtual screening due to its exceptional diversity achieved through the combination of LabNetwork’s global supplier network with WuXi’s scaffold design capabilities, its rapid synthetic accessibility with average delivery times of 4 to 8 weeks, and its built-in medicinal chemistry intelligence that helps reduce downstream attrition risks. This study utilizes the officially provided January 2025 version of the library, containing 29.1 billion compounds.

#### B.2 Conformer Generation

The molecules in both the Wuxi Galaxi and Enamine REAL libraries are provided as SMILES strings. As the DrugCLIP model requires three-dimensional structural coordinates for molecular encoding, a low-energy conformer was generated for each molecule using RDKit. The resulting atomic coordinates along with atom types were subsequently used as initial structural inputs to the DrugCLIP-based virtual screening pipeline.

#### B.3 Structural Clustering and Space Partitioning

The structural organization of the chemical space is critical to ensuring both the diversity of screening results and the feasibility of parallel processing. We implement a clustering strategy that partitions the hundred billion molecule library into multiple distinct subsets where each subset represents a specific chemical family or structural motif. This method prevents the output from being dominated by highly similar analogues by selecting top candidates from each cluster rather than just the global pool which allows for a balanced trade-off between scoring metrics and chemical diversity. Dividing the dataset into smaller manageable segments also facilitates massive parallelization as each partition can be searched independently to accelerate the overall screening timeline.

We employed a two-stage clustering-and-assignment strategy. First, compounds were randomly sampled to form a representative subset for initial clustering using K-Means. In the second stage, all compounds in the library were assigned to the nearest cluster center defined by Euclidean distance. Detailed analysis is provided in the Section E.1. When processing the full library, the 0.1% subset (94 million) was used to optimize 10,000 cluster centers. All compounds were then assigned to their nearest center, resulting in 10,000 partitions with sizes ranging from approximately 4 to 40 million compounds, with associated representations and metadata reorganized accordingly. Both K-Means clustering and distance calculation were performed using Faiss [27].

### C Implementation Details of GalaxyVS

#### C.1 Foundation: Contrastive Learning and Dense Retrieval

The development of ultra-large virtual screening begins with the choice of a foundational algorithm. Although physics-based free energy perturbation provides the most accurate estimation of binding affinity, its computational cost makes it impractical at trillion-scale. Traditional docking offers a more affordable alternative but still struggles when faced with hundreds of billions of candidates. Recent advances introduced DrugCLIP, a contrastive learning framework that aligns protein pocket and ligand representations to identify likely binders with high efficiency. To enable DrugCLIP-based search on an ultra-large library, the entire small-molecule collection is embedded into vector representations and stored on disk, resulting in approximately two hundred terabytes of data.

##### C.1.1 The DrugCLIP Algorithm

The DrugCLIP model uses two distinct Uni-Mol based encoders, one for molecules and one for protein pockets, which are aligned through contrastive learning. Each encoder is a transformer that processes 3D atomic features. The molecule encoder incorporates pretrained Uni-Mol weights, while the pocket encoder is pretrained via contrastive distillation on the ProFSA dataset to align with it. The model is optimized with a contrastive objective that brings the embeddings of matching protein-ligand pairs closer together in the representation space while pushing non-matching pairs apart. This is achieved using a batch softmax loss composed of two symmetric components: one for retrieving the correct ligand given a protein pocket:

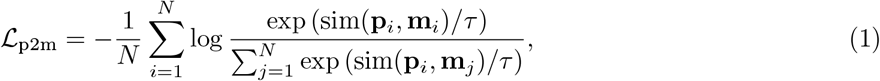

and another for retrieving the correct pocket given a ligand, with cosine similarity as the comparison metric:

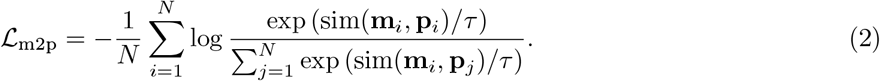

The total loss is the sum of these two terms:

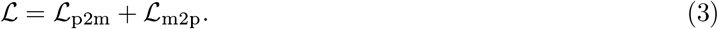

##### C.1.2 Dense Retrieval Framework

A key advantage of the DrugCLIP is its two completely separate encoders, which enable a highly efficient dense retrieval framework. In this approach, all molecules in a screening library are pre-encoded into vector representations offline and stored. During each new virtual screening run, only the target protein pocket requires on-the-fly encoding. Candidate molecules are then ranked by computing the cosine similarity between the single pocket embedding and all pre-computed molecule embeddings. This dense retrieval method can be accelerated further by dedicated search systems. For this work, we employed the PipeANN framework to optimize this large-scale similarity search, which is introduced in E.3.

To ensure robust and stable screening results, we use an ensemble of six models derived from cross-validation. During screening, the pre-encoded molecular representations from all six models and the newly encoded pocket embeddings from the same ensemble are used to generate predictions.

The scores are normalized using an adjusted robust z-score, which accounts for variations in score distributions when multiple pocket conformations are processed for a single target. The final score for each molecule is determined by selecting the maximum normalized score across all relevant pockets. The normalized score is called as adjusted robust z-score:

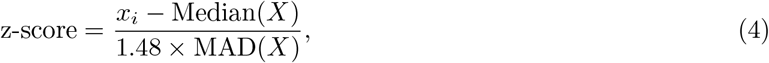

where

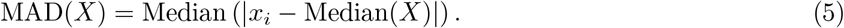

#### C.2 Hardware Adaptation and Large-Scale Molecular Encoding

We adapted the DrugCLIP workflow to a heterogeneous high-performance computing environment by replacing CUDA-based execution with custom accelerator backends integrated into a PyTorch-compatible framework (YH-Torch). Computational kernels were re-implemented as device-specific operators and registered through a unified operator interface, enabling transparent dispatch during execution while preserving the PyTorch programming model.

To improve efficiency for Transformer-based architectures, we implemented a fused multi-head attention operator that combines linear projections, matrix multiplications, softmax, and dropout into a single execution kernel. This reduces kernel launch overhead and improves on-device data reuse by minimizing intermediate memory transfers.

To ensure stable large-scale execution, we designed a node-aware task allocation strategy that assigns independent workloads to individual accelerator devices and partitions large datasets across jobs. Fault-tolerant execution and re-submission mechanisms were used to handle task-level failures and maintain throughput during long-running encoding jobs.

#### C.3 Vector Indexing

To ensure broad accessibility, we leverage PipeANN to support vector search directly against 100-billion-scale indices. PipeANN uses the same index structure as DiskANN, as shown in Figure 8. Vectors are organized as a directed graph, where each vector represents a node and is connected by edges. The index file is stored on disk to significantly reduce memory footprint, enabling ultra-large-scale screening on a limited number of standard nodes. To support higher performance, PQ-compressed vectors are stored in memory to accelerate access. Index traversal follows a deviated best-first search algorithm, which repeatedly executes 4KB random reads to access the current nearest vectors and its neighbor IDs to the query vector until convergence.

**Figure 8.**
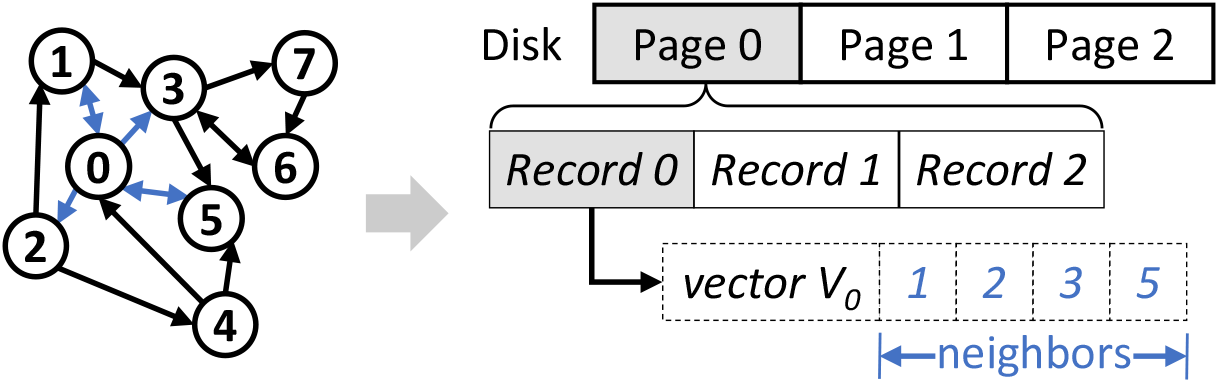
On-disk index layout of PipeANN.

In our cluster, the index files are stored in a Lustre parallel file system backed by hard disk drives (HDDs). While random reads suffer from high latency, PipeANN overlaps compute and disk I/O and uses an adaptive pipeline with a maximum I/O depth of 32 to hide part of the latency and utilize the parallelism of multiple HDDs. In our search scenario with a large top-*k*, the speculated I/Os in PipeANN lead to low latency with minimal I/O waste. This disk-native approach provides the foundation for the *Accessible Mode*. For massive parallel tasks, this architecture is further accelerated via a partitioning and preloading strategy in the *Extreme Mode* as detailed in the Section 2.3.

In our configuration, vectors are partitioned for parallel graph building and searching, where each partition contains approximately one million vectors (∼ 100,000 partitions in total). For most partitions, we set the maximum out-degree in the graph (*R*) to 64. In some hard partitions, search accuracy fails to be ensured using *R* = 64 (due to weak graph connectivity), so we increase their *R* to 128. The candidate pool size (*L*, larger values improve accuracy at the cost of higher latency) is set to 100 by default and to 192 for large *R* during index building, and to 5 × top-*K* during searching. We use 32 bytes for each PQ-compressed vector, which results in a memory-to-disk ratio of 1:128 (*<* 10TB memory in the cluster).

#### C.4 High-Performance In-Memory Retrieval

Genomic-scale virtual screening introduces significant challenges in simultaneously accommodating large molecular embedding databases and high query concurrency within limited aggregate memory. To address this, we adopt a distributed in-memory retrieval architecture on a supercomputing cluster.

Molecular embeddings are partitioned into shards and distributed across compute nodes, where each node loads its assigned partitions into local memory for graph-based search. This design preserves the efficiency of conventional in-memory vector retrieval while remaining compatible with the index structure used in the disk-based accessible search mode.

To support large-scale execution, data loading is carefully staged to avoid contention on shared storage, after which both query and database embeddings remain resident in memory throughout the search phase. This eliminates runtime I/O during retrieval and enables efficient parallel execution across tens of thousands of compute nodes.

The system supports partitioned execution with centralized aggregation of results following distributed searches.

#### C.5 Task Scheduling

Executing calculations on such a massive scale transforms low probability hardware anomalies into inevitable operational challenges. We deploy a stable and robust task scheduling system designed to manage the complex workflow and handle unexpected events such as computing node failures automatically. Ensuring the reliability of the scheduling process is essential for maintaining continuous operation and guaranteeing that all screening tasks reach completion without manual intervention.

This work involves a massive number of jobs. Each job requires a significant number of computing resources to finish within a reasonable amount of time, which can be executed *via* parallel processes with multi-threads. We designed a hybrid high-throughput computing (HTC) task management system in order to optimize the overall time to solution of all calculations, as shown in Figure 9. The systems perform dynamic job allocation by continuously fetching idle nodes from a constantly updated list of available compute nodes to perform unfinished jobs. Each job is characterized with its resource requirements. If a job/node failure was detected, then the job would be allocated another idle node. A job would be terminated and removed if it could not be finished after 3 trials, where the problem is most likely to be caused by the physics model rather than a hardware problem. Each node is released to the idle list once the assigned job is completed successfully, which allows execution of remaining jobs. If the number of idle nodes exceeds the requirement of remaining jobs, extra nodes will be released. The dynamically maintained queues and lists of jobs and nodes form an execution loop until all jobs are processed.

**Figure 9.**
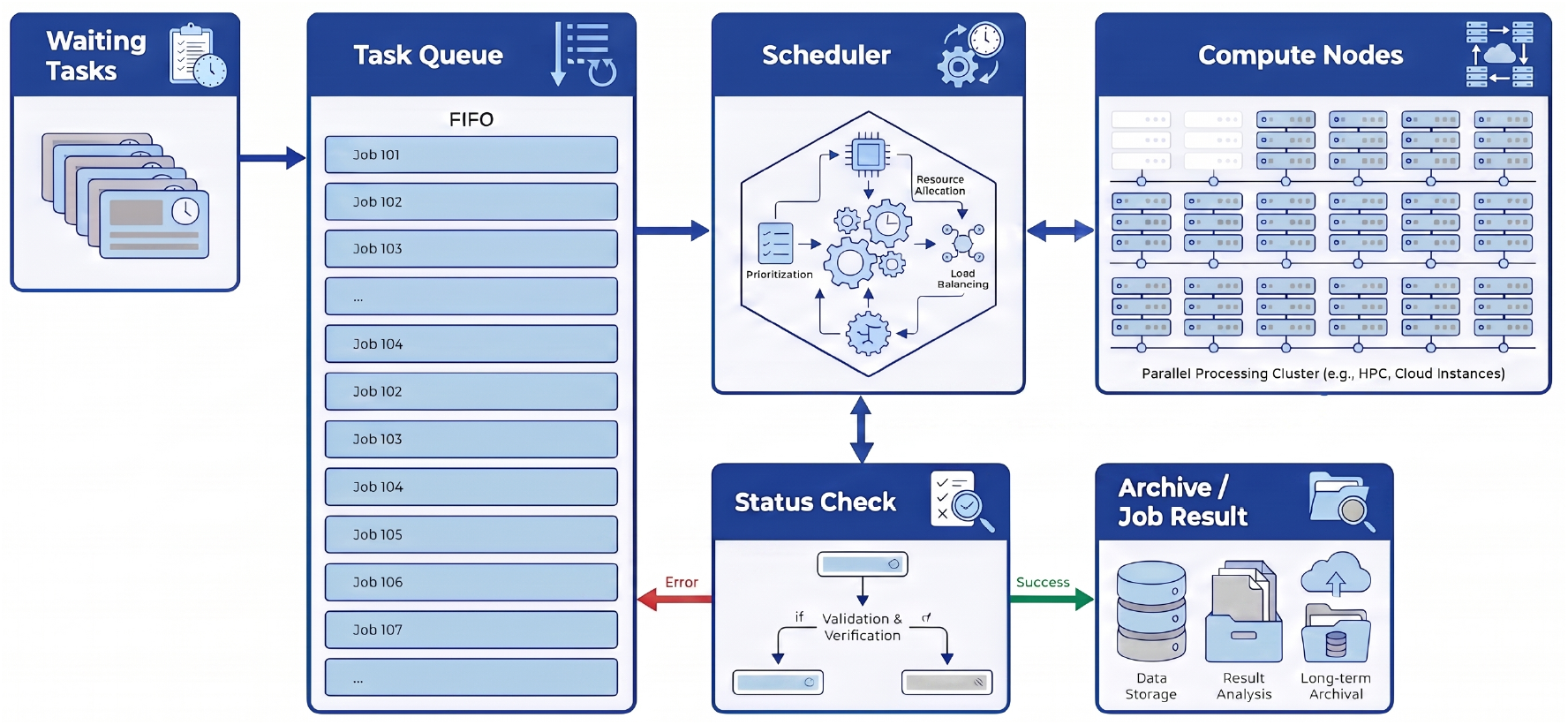
A hybrid high-throughput computing task management system.

When the management system checked that all simulation tasks were completed, the system should check whether each job is successfully finished.

#### C.6 Candidate Selection and Affinity-based Re-ranking

##### C.6.1 Property-and-rule-based Filtering

To preserve flexibility, the 100B library was not pre-filtered during data preprocessing or DiskANN index construction. Instead, filtering was applied to each retrieved and enriched subset individually, as filtering criteria may vary across targets and design objectives. For the multi-species genome-wide database construction, as well as diversity and predicted affinity evaluations across multiple targets, we adopted the same filtering rules used in DrugCLIP [11, 12]. These filters served as a basic quality control procedure. For single-target ABFE and experimental evaluation, we applied a looser rule set (Table 4) to better reflect realistic target-specific optimization scenarios.

**Table 4.** Physicochemical properties and structural alert limitations for molecular filtering.

| Property | Limitation |
| --- | --- |
| Molecular weight | [150, 550] |
| Number of rings | [1, 7] |
| Number of H-bond donors | [0, 6] |
| Number of H-bond acceptors | [0, 12] |
| ClogP | [-3, 5] |
| Topological polar surface area (TPSA) | [0, 200] |
| Number of rotatable bonds | [0, 12] |
| Number of aromatic rings | [1, 7] |
| Max size of ring | [3, 8] |
| Number of isomers | [1, 8] |
| Fraction of N or O | [0.001, 0.4] |
| Fraction of heteroatoms | [0.001, 0.5] |
| Number of contiguous rotatable bonds | [0, 5] |
| Number of contiguous non-ring bonds | [0, 6] |
| Number of contiguous rotatable bonds in scaffold | [0, 5] |
| Allowed atom types | {H, C, N, O, F, Cl, Br, I, S, P} |
| No matching structural alert catalogs | PAINS, ZINC |
| No matching patterns | Multi-ether-ester ([#6] - [#8, #16; !a] - [#6], count $\geq 4$)<br>Di-guanidine ([#7] ~ [#6] (~ [#7]) ~ [#7] ~ [#6] (~ [#7]) ~ [#7])<br>Nitro ([#7] (~ [#8]) ~ [#8]) |

##### C.6.2 Diversity Factor

After retrieving the top 0.01% of the 100B library, the resulting ∼ 10M focused set can still impose substantial computational overhead for downstream ranking methods, especially on small clusters or single-node setups. Directly selecting the top-scored compounds, especially when the selection size is small, can substantially reduce the chemical diversity of the candidate set. To address this, GalaxyVS incorporates a diversity control mechanism that explicitly regulates how many predefined clusters are represented when further selecting top-ranked compounds.

Specifically, we adopt a two-stage selection procedure. In the first stage, the top-ranked fraction *r* is independently selected from each cluster partition. In the second stage, the final top-*k* compounds are chosen from the union of all retained candidates. As *r* increases, the number of clusters represented in the final top-k set, denoted by *l*, increases monotonically from a minimum value up to min(*k*, 10,000).

Given a target top-*k* and a desired *l*, we use binary search to efficiently determine the optimal per-cluster retention ratio *r*. This procedure requires only a single global sorting of the ∼ 10M focused library, followed by iterative grouped slicing and merging are performed to count *l*, and can be completed within minutes in practice.

##### C.6.3 Affinity-based Re-ranking with AlphaRank

The initial retrieval stage effectively enriches the pool of potential binders but does not strictly order them according to biological affinity. To enhance the reliability and reference value of the final selection we incorporate a re-ranking stage utilizing a precise affinity prediction model. We employ AlphaRank which is built upon the AlphaFold3 backbone to evaluate and sort the enriched candidates based on their predicted interaction strength. This multi-stage pipeline condenses the screening of a hundred billion molecules for single or multiple targets into an efficient hourly workflow that significantly elevates the practicality of ultralarge-scale virtual screening.

### D Experimental Setup and Evaluation Protocols

#### D.1 Virtual Screening Benchmarks

Virtual screening is a critical component of early-stage drug discovery, aiming to identify a small number of structurally diverse chemical series with modest activity and favorable properties under limited experimental testing budgets. A central challenge lies in selecting tens of promising and chemically diverse candidates from libraries containing millions of available compounds using computational methods.

Benchmark datasets such as DUD-E and LIT-PCBA evaluate virtual screening performance using libraries either consisted of known active compound paired with computationally identified decoys or generated from large-scale experimental screening data. These benchmarks typically assess enrichment factors at conventional cutoffs (e.g., top 1%), measuring how effectively a method enriches actives relative to the full screening background.

However, ultra-large-scale virtual screening over libraries on the order of 100 billion compounds introduces new evaluation challenges. In practice, only a vanishingly small fraction of the library can be explicitly retrieved and examined, rendering standard metrics such as EF_1%_ inapplicable, while exhaustive ranking required for metrics like BEDROC is computationally infeasible.

To address this, we adopt an approximate evaluation strategy in which known active compounds are assigned to specific library partitions based on their distances to cluster centroids. For each target, we first perform ANN search to retrieve the top *k*% compounds from the ultra-large library. Known actives belonging to the corresponding partitions are then inserted into the ranked list according to their scores, with lower-ranked compounds displaced beyond the top *k*%, enabling the computation of EF_*k*%_.

**DUD-E [28]** is a widely used benchmark for structure-based virtual screening. It comprises a curated set of 102 protein targets spanning diverse families, including kinases, proteases, other enzymes, nuclear receptors, and transmembrane targets, selected to represent common druggable target classes. For each target, a complex structure and a collection of known active ligands (avg. 224) are provided, together with a large set of property-matched decoys (avg. 62 × actives) designed to mimic the physicochemical characteristics of the actives while being structurally distinct. While the decoys are computationally generated and may not fully reflect real screening libraries, the dataset remains valuable for assessing enrichment behavior across heterogeneous target classes.

**LIT-PCBA [29]** is another common used benchmark derived from experimentally high throughput screening data, designed to reduce chemical biases introduced by artificial decoys. It consists of 14 protein targets (one comprises both agonists and antagonists), 7,761 active compound and 382,674 inactives selected from high-confidence assay outcomes, with at least one complex PDB template. Unlike decoy-based benchmarks, LIT-PCBA directly reflects screening scenarios where inactives originate from real compounds tested experimentally. By relying on assay-derived inactives rather than synthetic decoys, LIT-PCBA presents a more realistic and challenging evaluation setting for virtual screening methods, but the active-to-inactive ratio varies substantially across targets.

#### D.2 Screening Metrics at the 100-Billion Scale

Given the prohibitive computational cost of obtaining a full global ranking over the entire library, metrics such as AUROC or BEDROC become infeasible at this scale. We therefore evaluate performance using enrichment factors at small cutoffs, which are more consistent with realistic screening settings. The enrichment factor is computed as:

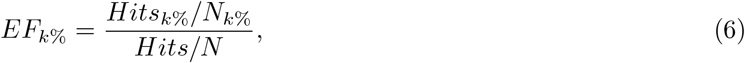

where *Hits*_*k*%_ denotes the number of active compounds retrieved within the top *k*% of the ranked list, *N*_*k*%_ is the total number of compounds in the top *k*%, and *Hits/N* corresponds to the overall positive rate of the screening library.

#### D.3 Diversity Assessment Metrics

We assess the diversity of enriched chemical sets from multiple complementary perspectives, including overall molecular similarity, as well as the abundance of unique scaffolds and fragments. All chem-informatics analyses were performed using RDKit [16], except for clustering, which was performed using Schrödinger suite.

##### Unique clusters

Clusters are constructed using the Leader–Follower clustering algorithm (ECFP 1024-bit, threshold = 0.85), with compounds processed in descending order of score.

##### Unique scaffold count

Scaffold diversity is quantified by counting unique Bemis–Murcko scaffolds [30] extracted from the molecules in the set.

##### Unique atomic environment

Atomic environment diversity is measured by enumerating unique atom-centered fragments defined by Morgan fingerprint environments [31] up to a radius of four bonds by covalent connectivity (4-hop neighborhoods). Specifically, each atom is first represented by a set of features mapped to an integer via a hash function to encode its atomic state. The integers of neighboring atoms are then combined with that of the center atom and hashed to generate new integers, with this process iteratively repeated to encode 1-hop, 2-hop, 3-hop, and up to 4-hop neighborhoods. Counting the unique integers across all atoms in the set serves as a proxy for the number of unique local atomic environments.

##### Unique BRICS fragments

Fragment diversity is further evaluated using BRICS [32] fragmentation, where molecules are decomposed by breaking synthetically plausible bonds using the BRICS.Decompose algorithm, and the number of unique BRICS fragments is reported.

#### D.4 Affinity Prediction and ABFEP Calculations

While virtual screening typically aims to identify compounds with modest binding activity, the ultra-large screening library contains orders of magnitude more compounds and is therefore expected to include substantially more target-active compounds and chemically distinct series than conventional commercial libraries. A key question is *whether this advantage in scale and diversity translates into improved hit quality*. To this end, we employ advanced affinity prediction models to compare the binding quality of equal-sized compound sets retrieved by GalaxyVS and by DrugCLIP on a common commercial library.

##### D.4.1 Screening Target Set

The screening target set was adapted from DUD-E [28], which provides a diverse collection of 102 well-characterized protein targets commonly used for structure-based virtual screening. The selected targets span a broad range of binding site architectures and functional classes, including 26 kinases, 15 proteases, 36 other enzymes, 11 nuclear receptors, 5 GPCRs, 2 ion channels, 2 CYP450s, and 5 additional targets.

Target sequences were obtained via the UniProt ID mapping web service. PDB IDs associated with DUD-E targets were mapped to UniProt IDs, and when multiple mappings were returned, the appropriate entry was manually selected based on the target name. For the INHA target, whose PDB structure (2H7L) has been marked obsolete and removed from the current PDB archive, we instead used its updated replacement ID (4TRJ).

##### D.4.2 Predicted Affinity Score

Given the prohibitive cost of experimentally determining binding affinity for large-scale candidate sets, we employed two advanced computational alternatives, Boltz-2 [17] and AlphaRank, for high-throughput evaluation. Both models share a core architectural premise: they utilize AlphaFold3 as a foundational backbone for structural feature extraction, followed by dedicated prediction heads for estimating binding affinity. Their training leverages large-scale bioactivity data from sources like ChEMBL and BindingDB. The key distinction lies in their specific learning objectives. AlphaRank formulates affinity prediction as a ranking task, optimized with a pairwise ranking loss to order molecules by binding strength without predicting absolute values. In contrast, Boltz-2 is trained to directly predict quantitative affinity scores alongside a binary binding classification. In terms of performance, both models offer accuracy comparable to far more computationally intensive methods like free-energy perturbation, while operating at a fraction of the cost. They demonstrate strong and complementary predictive capabilities across different protein targets. The affinity scores predicted by these models thus serve as a robust, efficient, and standardized computational metric for evaluating the potential binding strength of molecules identified in our virtual screening campaign.

##### D.4.3 ABFEP Preparation and Calculation

BRD4 binding regions from five crystal structures (PDB IDs: 5UF0, 5UEZ, 5UEW, 5UEU, and 5UEY) were used as queries for virtual screening. From the top 0.01% of results, 50,000 molecules representing 5,000 unique indices were retained. Following general physicochemical and structural filtering, candidates were clustered using a leader–follower algorithm based on ECFPs with a distance threshold of 0.85.

Cluster representatives were docked into the 5UF0 receptor. For each ligand, only the top docking pose (rather than the default 5) was retained for subsequent ABFE calculations [19]. As docking scores do not reliably rank poses for ABFE, this may not select the lowest free energy conformation and yields more conservative estimates. All other simulation parameters followed the default settings of the BRD4 demo system. In addition, the top 10 molecules ranked by docking score were included for docking pose quality control.

### E System Validation and Experimental Analysis

#### E.1 Comparison of Large-Scale Clustering Methods

Several clustering methods were evaluated for their ability to handle even this large subset, including the distance-based algorithm BitBirch [33], a combination of MinHashLSH [34] for approximate distance computation with Leader-Follower clustering, and K-Means (FAISS [13] implementation). Precomputed ECFP4 molecular fingerprints were used to represent compound structures during clustering.

For evaluation, we randomly sampled 0.01% of the full library, corresponding to 9.4 million compounds. From this subset, a further random 0.1% (9.4 thousand compounds) was selected to compute clustering centers.

Classical distance-based molecular clustering methods, including Butina [35] and the K-Means variant K-Medoids (since fingerprints are binary vectors), were used as baselines for first-stage clustering. Although these methods are computationally infeasible for clustering the full library, they provide useful references for assessing clustering quality.

We report the number of (non-singleton) clusters, the number of singletons, and the maximum cluster size for each method. In addition, clustering quality was evaluated using the Calinski–Harabasz Index [36] (CHI), Davies–Bouldin Index [37] (DBI), and Dunn Index [38] (DI). Given the large scale of the evaluation set, we adopted the iSIM [39] variants of these indices implemented within the BitBirch framework.

The results for both stages were provided in the Supplementary Material. In the clustering stage, distance-based methods produce clusters that are more compact and better separated, as reflected by higher CHI and DI, whereas K-Means achieve lower DBI, indicating more uniform inter-cluster similarity and more balanced cluster sizes. In the assignment stage, we chose K-Means clustering with Euclidean distance, which yields relatively uniform cluster sizes and the fastest runtime, with clustering metrics comparable to alternative methods.

#### E.2 Hardware Platform Consistency Validation

Numerical results can vary across different computational platforms. In addition to evaluating GalaxyVS’s performance via enrichment, affinity, and manual inspection, we assessed potential numerical differences and ranking consistency between the accelerator and the original GPU device (NVIDIA A100 80GB).

We first randomly sampled 1 million molecules and encoded them on both platforms. Element-wise differences were observed at the 4^th^–5^th^ decimal place, with a mean ± standard deviation of 6.86 × 10^−5^ ± 5.76 × 10^−5^ and a maximum deviation of 5.75 × 10^−3^. Next, we computed pocket representations for 744 conformers across 53 pocket groups. Considering the per-conformer ranking and intra-pocket group ranking (a total of 797 ranking pairs), Spearman’s correlation indicated agreementof at least 6 digits, with discrepancies in the top 1% (∼ 10k) ranging from 8–16 molecules. Furthermore, using 100 randomly sampled and normalized arrays as pocket representations, the Spearman correlation indicated agreement between 5–6 digits, and the top 1% differences were 6–26 molecules, corresponding to 0.1%–0.2% of the subset.

Overall, these results indicate that the accelerator and GPU produce minor numerical differences, which remain within a practically acceptable range.

#### E.3 Retrieval Precision of PipeANN

Building upon the dense retrieval framework, we evaluated the retrieval precision of PipeANN. As an Approximate Nearest Neighbor search engine, PipeANN achieves billion-scale retrieval efficiency by substituting exhaustive distance computations with heuristic proximity graph traversals. The inherent trade-off of this approximation is that the search path may occasionally converge to a local optimum, leading to minor precision errors where true global nearest neighbors are omitted.

For early-stage virtual screening pipelines, a minor omission rate below 5% is generally acceptable. This margin is tolerated because the initial dense retrieval serves as a coarse filter over an exceptionally vast chemical space, and downstream structure-based methods will subsequently perform rigorous evaluations on the retained candidates.

To quantify this precision and ensure the approximation remains within acceptable bounds, we evaluated the exact retrieval configurations used in our screening pipeline. During the actual search process, we query each separate index shard to retrieve the top 100 nearest neighbors using a search list size L set to 500 (5 × top-*K*). Under these exact parameters, we randomly selected three pre-built indices to assess their exact match overlap against exhaustive baseline computations. The results are summarized in Table 5. The system achieves an average recall rate of 97.67%, consistently remaining well within the acceptable error bounds. This confirms that the retrieval module maintains robust and highly accurate performance despite the use of approximate calculations.

**Table 5.** Retrieval precision of PipeANN across randomly selected indices.

| **L** | **Top K** | **Recall (%)** |
| --- | --- | --- |
| 500 | 100 | 98.00 |
| 500 | 100 | 97.00 |
| 500 | 100 | 98.00 |
| **Average** |  | **97.67** |

**Table 6.**
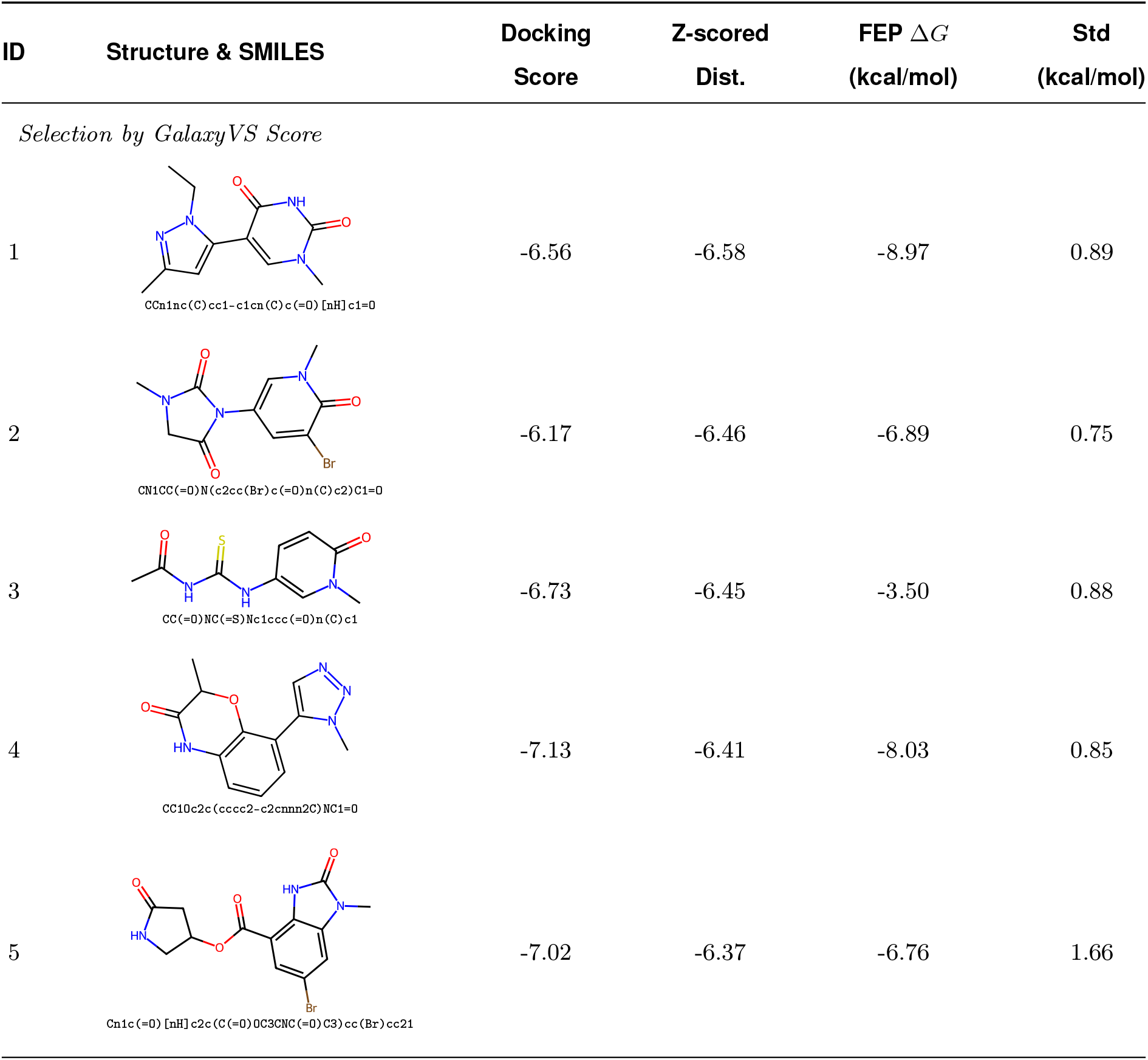
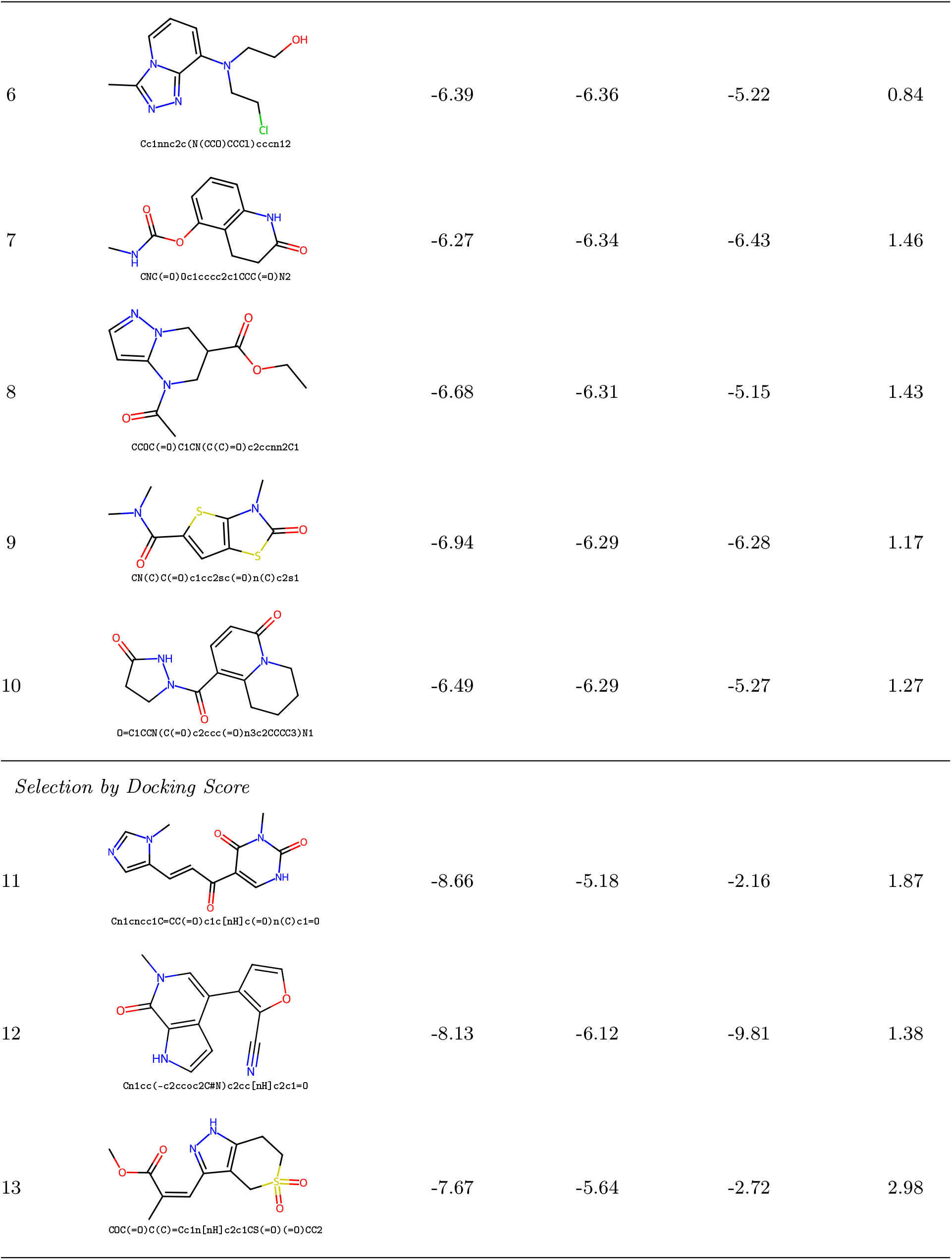
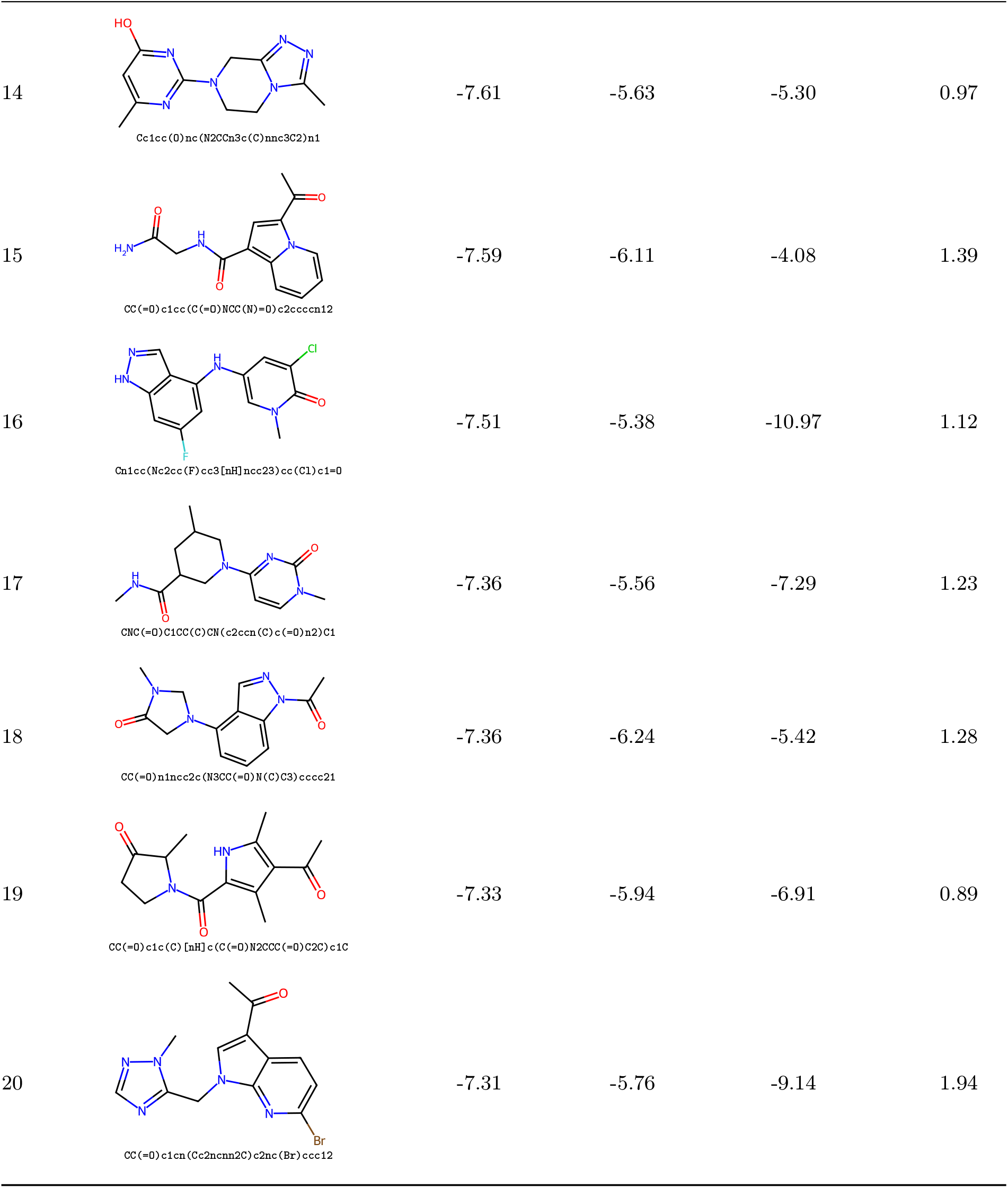
ABFEP validation results of candidates selected by GalaxyVS and Docking score. Values marked with “–” indicate a failure in converting the Tripos Mol2 file during ligand initialization.

**Table 7.** AlphaFold2 structural data for representative species.

| Species | Type | Total Structures |
| --- | --- | --- |
| Human (*Homo sapiens*) | Mammal | 23,586 |
| Mouse (*Mus musculus*) | Mammal | 21,452 |
| Budding yeast (*Saccharomyces cerevisiae*) | Fungus | 6,055 |
| Fruit fly (*Drosophila melanogaster*) | Insect | 13,461 |
| *E. coli* (*Escherichia coli*) | Prokaryote | 4,370 |
| *Arabidopsis* (*Arabidopsis thaliana*) | Plant | 27,402 |
| **Total** | - | **96,326** |

#### E.4 GalaxyVS achieves extreme enrichment under ultra-large-scale background

To assess whether GalaxyVS retains DrugCLIP’s enrichment power under an ultra-large compound background and a narrow retrieval regime, we evaluated enrichment at 0.01% using target and active compound data from common virtual screening benchmarks, including DUD-E [28] and LIT-PCBA [29]. As exhaustive scoring over the full 100-billion compound space is intractable, we adopted an approximate yet scalable evaluation protocol. Pocket and known active compound embeddings were precomputed on accelerator nodes. For each target, pocket embeddings were used to query clustered DiskANN [40] indices of the 100-billion compound library, retrieving the top 0.01% candidates ranked by L2 distance in the shared embedding space. The enrichment factor at 0.01% (EF_0.01%_) was computed with respect to the aggregated retrieved background across all partitions.

GalaxyVS achieves an EF_0.01%_ of 1594.3 on DUD-E, corresponding to a 1594-fold relative to the overall positive rate of the library, and an EF_0.01%_ of 297.6 on LIT-PCBA. These results indicate that GalaxyVS preserves strong enrichment capability even at an extreme retrieval depth of 0.01%, consistent with the trend observed on original benchmarks where EFs increase as the retained fraction decreases. Notably, the reduction in EF relative to the theoretical maximum is expected when replacing decoy-based or inactive-only backgrounds with a large-scale screening library, as the latter is more likely to contain previously unannotated active compounds.

#### E.5 Comparison of Selection Metrics within the Enriched Candidate Pool

As ABFEP initial conformations were derived from molecular docking, we assessed the relationship between docking scores and free energy estimates. To this end, we evaluated an additional 10 candidates selected by docking score from the same enriched and clustered pool. The comparison suggests a trade-off between calculated affinity and stability. Docking-based selection identifies candidates with lower apparent energy minima, for example Compound **16** (Δ*G* = -10.97 ± 1.12 kcal/mol), but introduces greater variability. Two such candidates (Compounds **11** and **13**) exhibit notably weak affinities (Δ*G* ≥ -3.00 kcal/mol), accompanied by large computational fluctuations (std up to 2.98 kcal/mol), indicating binding structural instability and partial pocket escape in explicit solvent. In contrast, GalaxyVS prioritizes molecules with more consistent and well-converged thermodynamic behavior over favorable static docking fits.

### F Curation of the GalaxyDB

#### F.1 Species Data for Virtual Screening

To evaluate the cross-species applicability of our ultra-large-scale virtual screening system, we processed the entire proteomes of multiple representative biological organisms. The structural data were sourced from the AlphaFold Protein Structure Database. Table 7 summarizes the original number of structures for each species prior to pocket identification and structural filtering. These raw counts provide a foundational reference for the comprehensive scale of the screening campaign.

#### F.2 Building a Representative Set for Database Curation

To balance enrichment quality and structural diversity, we performed an elbow analysis on the relationship between z-score thresholds and diversity factors across 102 DUD-E targets. Based on the median elbow point across targets, we selected a target diversity level of approximately 7,000 clusters. For each filtered enriched subset at the million-scale level per pocket, we retained 30,000 molecules while maximizing coverage of the 7,000 clusters under the additional constraint of z-score *<* −4.

To construct a representative and structurally diverse subset, the resulting 30,000 molecules were clustered using a Leader–Follower algorithm based on RDKit ECFP4 fingerprints (1024-bit). Cluster representatives were selected as the highest-ranked molecule within each cluster. The clustering threshold was defined as a Tanimoto similarity cutoff of 0.8.

#### F.3 Complex Generation through Ensemble Docking

Following retrieval and clustering, ensemble molecular docking was performed to generate candidate binding poses. This procedure complements retrieval-based implicit interaction representation with explicit geometric and physicochemical constraints.

Docking was conducted using AutoDock Vina (v1.2.5). Binding pockets were defined using ligand coordinates obtained from either the generation–sidechain-packing protocol or homologous structural alignment following DrugCLIP. Ligand conformers were generated with Open Babel, and both ligand and receptor structures were processed into docking-ready PDBQT files using Meeko.

For the Human dataset, docking was performed against 26,562 homologous or geometric pockets, represented by 187,715 pocket conformers in total (up to 10 conformers per pocket). Each pocket was docked with approximately 200 candidate ligands on average using an exhaustiveness of 16. For each ligand, the best-scoring pose across all available pocket conformers was retained for downstream analysis and database construction.

## Notes

https://galaxyvs.drugclip.com/

### Competing Interest Statement

The authors have declared no competing interest.

